# RNA-binding and HEPN-nuclease activation are decoupled in CRISPRCas13a

**DOI:** 10.1101/190603

**Authors:** Akshay Tambe, Alexandra East-Seletsky, Gavin J. Knott, Jennifer A. Doudna, Mitchell R. O’Connell

## Abstract

CRISPR-Cas13a enzymes are RNA-guided, RNA-activated ribonucleases. Their properties have been exploited as powerful tools for RNA detection, RNA imaging and RNA regulation. However, the relationship between target RNA binding and HEPN (higher-eukaryotes-and-prokaryotes nucleotide-binding)- domain nuclease activation is not well understood. Using sequencing experiments coupled with *in vitro* biochemistry, we find that Cas13a’s target RNA binding affinity and HEPN-nuclease activity are differentially affected by the number of and position of mismatches between the guide and target. We identify a central ‘binding seed’ where perfect base pairing is absolutely required for target binding, and a separate ‘nuclease switch’ where imperfect base-pairing results in tight binding but no HEPN-nuclease activation. These results demonstrate that the binding and cleavage activities of Cas13a are decoupled, highlighting a complex specificity landscape. Our findings underscore a need to consider the range of effects off-target recognition has on Cas13a’s RNA binding and cleavage behavior for RNA-targeting tool development.

## INTRODUCTION

Prokaryotic adaptive immune systems use CRISPRs (clustered regularly interspaced short palindromic repeats) and CRISPR associated (Cas) proteins for RNA-guided cleavage of foreign genetic elements (Mohanraju et al., 2016; Wright et al., 2016). Type VI CRISPR-Cas systems include a single protein, Cas13 (formerly C2c2), that when assembled with a CRISPR-RNA (crRNA) forms a crRNA-guided RNA-targeting effector complex (Abudayyeh et al., 2016; East-Seletsky et al., 2016; Konermann et al., 2018; Shmakov et al., 2015; Smargon et al., 2017; Yan et al., 2018). Cas13 proteins are classified into distinct subfamilies (Cas13a-d), and all Cas13 proteins studied to date possess two enzymatically distinct RNase activities that are required for optimal interference (Abudayyeh et al., 2016; East-Seletsky et al., 2016; Konermann et al., 2018; Smargon et al., 2017; Yan et al., 2018). First, upon binding a precursor-crRNA (pre-crRNA), Cas13 cleaves within the crRNA direct repeat in a pre-crRNA array to form mature Cas13-crRNA complexes (East-Seletsky et al., 2016; East-Seletsky et al., 2017; Konermann et al., 2018; Smargon et al., 2017; Yan et al., 2018). Second, binding of an RNA target complementary to the crRNA (henceforth referred to as an activator-RNA) triggers Cas13 to cleave RNA non-specifically by activating the enzyme’s two HEPN-domains to form a single composite RNase active site (Abudayyeh et al., 2016; East-Seletsky et al., 2016; Konermann et al., 2018; Liu et al., 2017a; Liu et al., 2017b; Smargon et al., 2017; Yan et al., 2018). This HEPN-activated conformation of Cas13 is a general nuclease, which is capable of cleaving either the RNA molecule that it was activated by (*cis* cleavage), or any other RNA molecule that it happens to encounter (*trans* / collateral cleavage). All Cas13 proteins characterized to date exhibit these behaviors (Abudayyeh et al., 2016; East-Seletsky et al., 2016; East-Seletsky et al., 2017; Gootenberg et al., 2018; Gootenberg et al., 2017; Konermann et al., 2018; Smargon et al., 2017; Yan et al., 2018).

The RNA-activated RNA cleavage behavior of Cas13 provides a mechanism for RNA detection and diagnostic applications (East-Seletsky et al., 2016; Gootenberg et al., 2018; Gootenberg et al., 2017), as well as RNA-targeting in heterologous systems (Abudayyeh et al., 2017; Aman et al., 2018; Cox et al., 2017; Konermann et al., 2018). Versions of Cas13 where the HEPN-domains have been catalytically inactivated (Abudayyeh et al., 2016; East-Seletsky et al., 2016; Konermann et al., 2018; Smargon et al., 2017; Yan et al., 2018) (deactivated Cas13; dCas13) have shown to be useful as programmable RNA binding proteins for RNA imaging (Abudayyeh et al., 2017), RNA editing (Cox et al., 2017) and regulating specific transcripts RNA in other ways (e.g. regulating pre-mRNA splicing (Konermann et al., 2018)). However, the relationship between crRNA: activator-RNA sequence complementarity and Cas13 binding specificity and HEPN-nuclease activation have not been determined, and based on the behavior exhibited by other CRISPR-Cas systems (for review see Jackson et al. (2017)), it is likely that this relationship is complex.

We used a combination of library-based high-throughput RNA-binding assays and biochemical experiments to interrogate the effects of mismatches on both activities of Cas13a from *Leptotrichia buccalis* (Lbu), enabling us to determine the relationship between activator-RNA recognition specificity and HEPN-nuclease activation. We find that both the number and distribution of mismatches between the crRNA and activator-RNA affect Lbu-Cas13a’s binding affinity and HEPN-nuclease activity. Mismatches in the middle ‘seed’ region (positions 9-12) of the crRNA-spacer result in the largest defect in binding affinity. In contrast, mismatches elsewhere in the crRNA result in only small reductions of binding affinity. Surprisingly, although targets with mismatches at positions 5-8 of the crRNA-spacer bind with moderate to high affinity, these targets fail to activate Lbu-Cas13a’s HEPN-nuclease activity. Conversely, a separate subset of mismatched, weakly-bound activator-RNAs were able to maximally activate Cas13a’s ribonuclease activity. These results show that activator-RNA binding and HEPN-nuclease activation can be decoupled in Lbu-Cas13a, revealing a mechanism that is not reflected by simple activator-RNA binding affinity but instead a complex specificity landscape that enables Lbu-Cas13a to distinguish between complementary and mismatched RNA transcripts prior to crRNA-guided RNase activation, all while allowing for relaxed RNA sequence specificity. Together, we show that the accuracy of activator-RNA binding and RNA-activated RNA cleavage by Cas13a enzymes are different, which has implications for understanding the role of Cas13a in bacterial immunity, as well as the development of Cas13a as a high-fidelity RNA-targeting tool.

## RESULTS

### High-throughput profiling of Lbu-Cas13a activator-RNA binding preferences

To determine the effects of individual crRNA: activator-RNA mismatches on the binding interaction with Lbu-Cas13a, we designed an *in vitro* library-based assay to assess binding mismatch tolerances and the effects of sequence variation outside the Cas13a: crRNA binding site. Our experimental design included two different sequence libraries that were mutagenized in the same manner (**Fig. 1A; see Methods)**, but only one crRNA was used in any given binding experiment, allowing us to distinguish between weak crRNA-dependent target binding and crRNA-independent nonspecific binding. The libraries were mixed at a 1:1 ratio and incubated separately with two concentrations of biotinylated HEPN-nuclease-deactivated Lbu-Cas13a (Lbu-dCas13a) protein (**see Methods & Fig. S1A**) complexed with crRNA. After incubation, bound RNAs were eluted and subjected to Illumina-based sequencing (**Fig. 1B; Table S1**). Controls were conducted to determine input library distribution (**Fig. 1C**) and the effect of non-specific Lbu-dCas13a background binding (**Fig. S1B-C).**

**Fig. 1.**
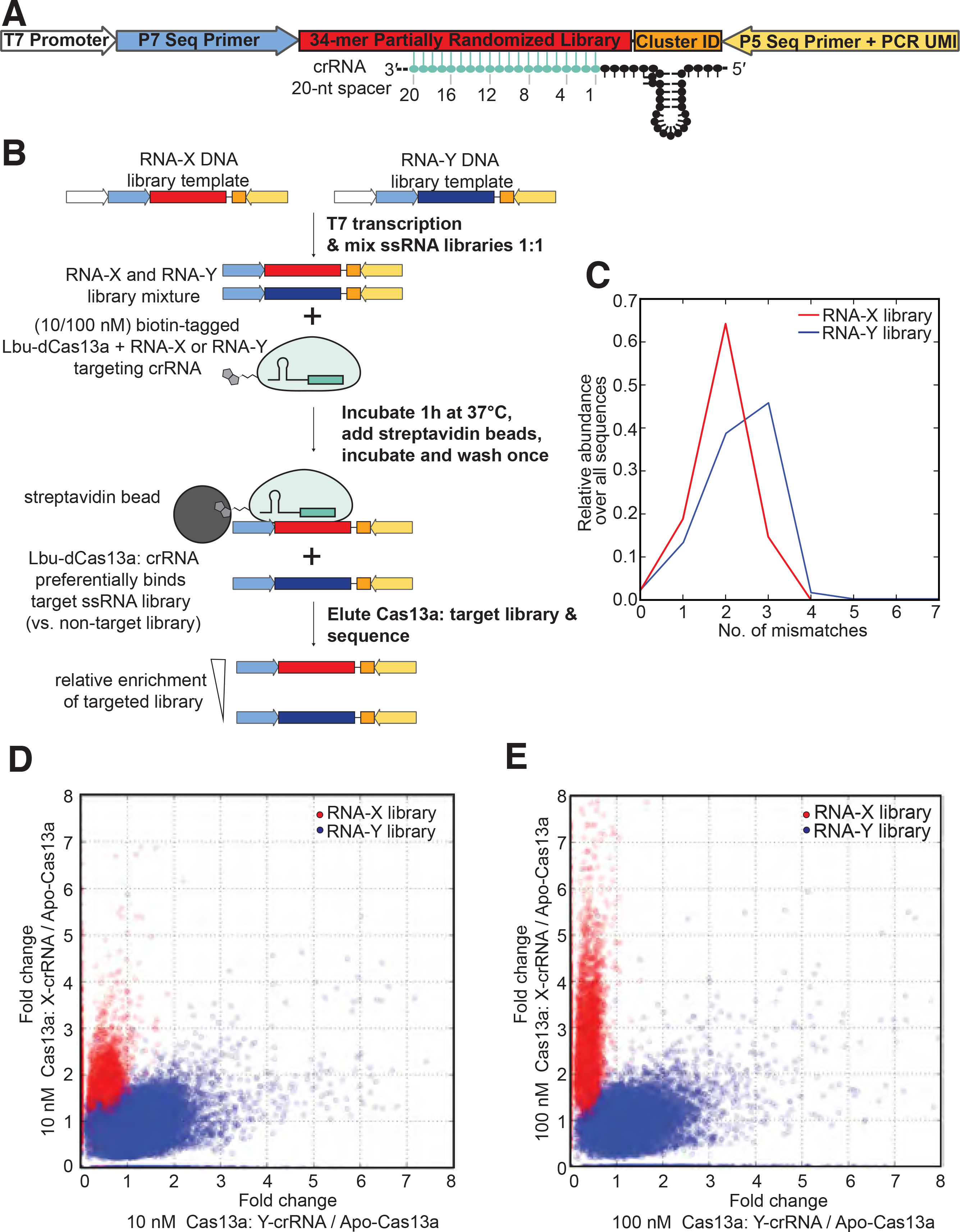
Development of High-throughput RNA mismatch profiling to determine activator-RNA binding preferences for Lbu-Cas13a. (**A**) Schematic of the high-throughput RNA mismatch profiling library design. T7 promoter: for producing RNA via in vitro transcription. This sequence lost upon *in vitro* transcription of the DNA template (see Methods). P7 Seq Primer: binding site for the P7 Illumina sequencing primer that includes an index for multiplexing library samples. 34-mer Partially Randomized Library: either RNA-X or -Y target sequence with 76% frequency of on-target nucleotides and 8% frequency of each other nt. at each position. 20nt-spacer crRNA binding site positon is shown. Cluster ID: 6-mer randomized region to ensure robust Illumina sequence cluster calling. P5 Seq Primer + PCR UMI: a modified P5 Illumina sequencing primer binding site that incorporates a 13-mer randomized ‘unique molecular identifier’ (UMI) within the index region to filter out PCR duplicates downstream. (**B**) Schematic of the high-throughput RNA mismatch profiling pulldown workflow. (**C**) empirical mismatch distribution profile for the input X and Y activator-RNA libraries (prior to Cas13a binding). (**D**) A scatter plot displaying fold-change in abundance of each RNA-X and -Y library members between samples with 10 nM crRNA-bound Cas13a (vs. crRNA-unbound apo-Cas13a) programmed to target RNA-X or -Y sequences. (**E**) A scatter plot displaying fold-change in abundance of each RNA-X and -Y library members between samples with 100 nM crRNA-bound Cas13a (vs. crRNA-unbound apo-Cas13a) programmed to target RNA-X or -Y sequences.

We developed a computational pipeline to analyze our sequencing data (**see Methods**) that first removes PCR duplicates (by counting unique molecular identifier sequences), and then counts the number of observations for each library variant in a given sample. For each variant, we identified which input sequence it was derived from by Hamming distance, and recorded its relative abundance in a sample. After applying a threshold to filter out low-abundance library members, data from the remaining ~18,000 unique sequences were highly reproducible across replicates, with Pearson R >= 0.9 for all but one case (**Table S1**). We identified the crRNA-dependent signal from our data by computing the fold-change in abundance between pulldowns with crRNA-bound Lbu-dCas13a and crRNA-unbound apo-Lbu-dCas13a. A scatter plot of this fold-change for each of the different crRNA containing samples (X and Y) showed that target sequences that were strongly enriched by Lbu-dCas13a: crRNA-X were depleted by the Lbu-dCas13a: crRNA-Y, and vice versa (**Fig. 1D, E**).

Unsurprisingly, in each case, the enriched and depleted sequences corresponded to the on-target and off-target activator-RNA libraries, respectively. This clear signal of orthogonally indicated that the sequence enrichment and depletion behavior we observed is crRNA-dependent. Interestingly, we observed that this orthogonal signal was significantly diminished when comparing sequence enrichment by Lbu-dCas13a: crRNA relative to the input library, rather than relative to the crRNA-unbound apo-Lbu-dCas13a sample. (**Fig. S1B-C**). This implies that apo-Lbu-dCas13a protein has a non-negligible affinity for RNA that is sequence and/or structure dependent. Although we control for this in the analysis presented here, this property might need to be considered when using this Cas13a as an RNA-targeting tool.

### High-throughput profiling identifies activator-RNA mismatch sensitivity hotspots

We calculated the relative enrichment of target RNA library members with exactly one mismatch to each crRNA-guided Lbu-dCas13a sample, revealing several interesting observations **(Fig. 2A-B; Fig. S2)**. First, as expected, the magnitude of enrichment for each single-mismatch member within each targeted library was highly dependent on the position of the single-mismatch within the crRNA-targeting region, but independent of which nucleotide was present on the target at the mismatched site. Second, we observed that single-mismatch members were significantly less enriched (see Methods for significance testing procedure) when their mismatches occurred within a contiguous stretch of nucleotides in guide-RNA positions 9-14 compared to all other positions with the effect most pronounced at high Lbu-dCas13a: crRNA concentration (**Fig. 2A; Fig. S2).** This suggests that single base pair mismatches in positions 9-14 can significantly decrease Lbu-dCas13a’s affinity for activator-RNA. Conversely, library members with single base pair mismatches outside these regions (i.e. in guide-RNA sequence positions 1-5 & 15-20) are enriched as well as or even greater than the perfectly complementary target RNA, indicating that single mismatches in these positions in the target RNA: crRNA duplex have little to no effect on the overall binding affinity relative to a perfectly complementary target. Unexpectedly in some cases, they may even promote tighter binding than observed for perfectly complementary activator-RNA (**Fig. 2A**).

**Fig. 2.**
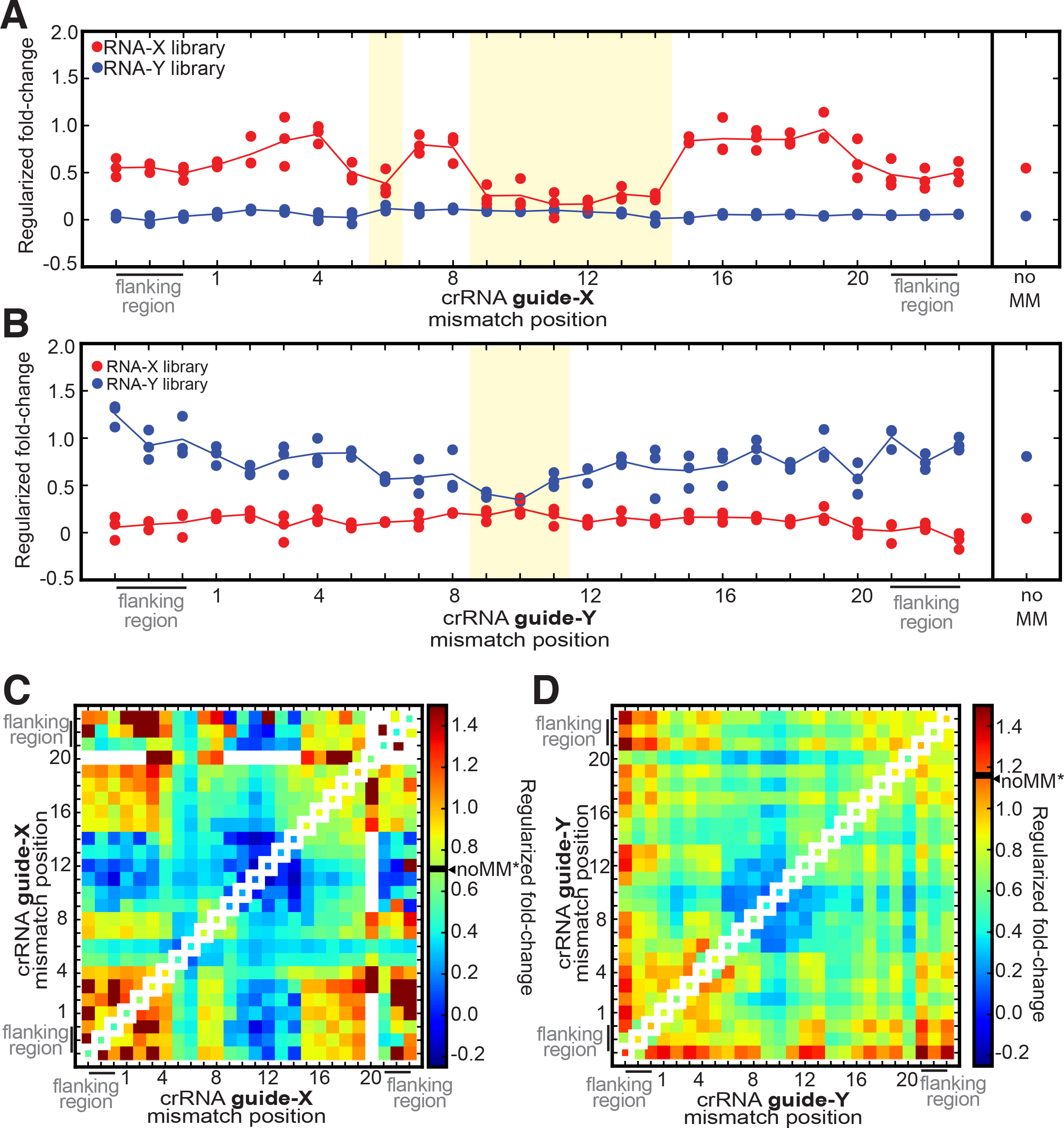
dCas13a high-throughput RNA mismatch profiling confirms binding mismatch sensitivity hotspots for single- and double-mismatched activator-RNAs. Regularized fold-change enrichment analysis (relative to Apo-dCas13a) for activator-RNA members of both libraries that contain only a single mismatch across the shown spacer-targeting and flanking regions when (**A**) targeting X activator-RNA library with 100 nM dCas13a: crRNA-X, (**B**) targeting Y activator-RNA library with 100 nM dCas13a: crRNA-Y. In each plot, each individual point in a position represents the average regularized fold-change value for each unique mismatch combination possible in that position (from n=3 experiments). The solid line represents as an average fold-enrichment (of all mismatches combinations) in that position. The fold-change enrichment of the perfectly complementary (noMM) targets in each experiment are show on the right of each plot. Permutation tests (100,000 permutations) were carried out to determine whether statistical significant differences/similarities between mean fold changes of each library exist at each mismatch position. Positions shaded in yellow indicate where statistically significant (p < 0.05) lack of enrichment between guide-complementary and guide-noncomplementary sequences was observed. See Methods for regularization and permutation test procedure. See **Fig. S2** for 100 nM Cas13a: crRNA data plotted as individual data points from three experiments, as well as individual and averaged 10nM Cas13a: crRNA data. Regularized fold-change enrichment analysis (relative to Apo-dCas13a) for activator-RNA members of both libraries for all pairs of mismatches across the shown spacer-targeting and flanking regions when **©** targeting X activator-RNA library with 100 nM dCas13a: crRNA-X, and **(D)** targeting Y activator-RNA library with 100 nM dCas13a: crRNA-Y. For comparison, single mismatch data is plotted along the diagonal axis and the fold-change value for a perfectly complementary target is indicated on the heat-map scale bar (noMM*).

A qualitatively similar and statistically significant effect is seen when the RNA-Y library is targeted by Lbu-dCas13a: crRNA-Y, where single-mismatches in positions 9-11 of the crRNA-spacer are also significantly less tolerated **(Fig. 2B; Fig. S2**). The difference observed in the overall magnitude of mismatch sensitivity when comparing RNA-X and RNA-Y target libraries could be due to crRNA and target RNA sequence composition, secondary structure propensity of the crRNA and/or target RNA, and the overall efficiency of the sample recovery.

We also calculated the relative enrichment of activator-RNA library members with two mismatches, and visualized the resulting data using a heatmap. This analysis has the potential to show interactions between different positions of mismatches: it can uncover combinations of two mismatches that act constructively to increase the binding defect (when compared to either singly mismatched target), or pairs of mismatches whose effects compensate for each other leading to tighter binding than that observed for either of the singly mismatched targets alone. In both the RNA-X and RNA-Y sequences, we clearly identify a central mismatch sensitive region (positions 9-14 for RNA-X, 9-11 for RNA-Y) where pairs of mismatches show even weaker binding affinity (when compared to single mismatches) (**Fig. 2C-D**; **Fig. S3).** Surprisingly, although single mismatches at positions 5-8 also show moderate binding defects, we find that mismatches in these positions are able to somewhat compensate for the loss of binding affinity caused by mismatches at positions 9-12 (**Fig. 2C).** These observations again suggest that the effect of individual mismatches on binding may not be additive when in combination, hinting that complex contributions from individual and sets of base-pairing interactions influence overall affinity for activator-RNAs.

Finally, it should be noted that we found little evidence that sequence elements outside the crRNA: target RNA duplex (termed the protospacer-flanking motif or PFS (Abudayyeh et al., 2016; Liu et al., 2017b)) contribute to crRNA-guided Lbu-dCas13a binding to target sequences (see ‘flanking regions’ in **Fig. 2C-D, Fig. S3**). Thus, at least for Lbu-Cas13a, a PFS isn’t required for optimal activator-RNA binding.

### Biochemical validation of Cas13 binding profiles

We independently validated the mismatch-sensitive profiles identified through high-throughput binding assay, by employing a fluorescence anisotropy assay to measure target RNA binding to Lbu-dCas13a. We paired activator-RNAs containing sets of four contiguous nucleotide mismatches with an otherwise complementary crRNA (**Fig. 3A**), tiled across the 20-nt. guide sequence **(Fig. 3B).** For three out of five of these activator-RNAs (activator-RNAs g1-4MM, g13-16MM, g17-20MM), the affinity of the crRNA-Cas13a interaction decreased by 6- to 8- fold compared to the interaction with a fully complementary RNA (noMM; **Fig. 3B, C).** Importantly, as expected from the high-throughput binding experiment, mismatches in nucleotide positions 9-12 of the crRNA spacer (activator-RNA g9-12MM) led to the largest binding defect, a 47-fold decrease in binding affinity compared to a fully complementary ssRNA.

**Fig. 3.**
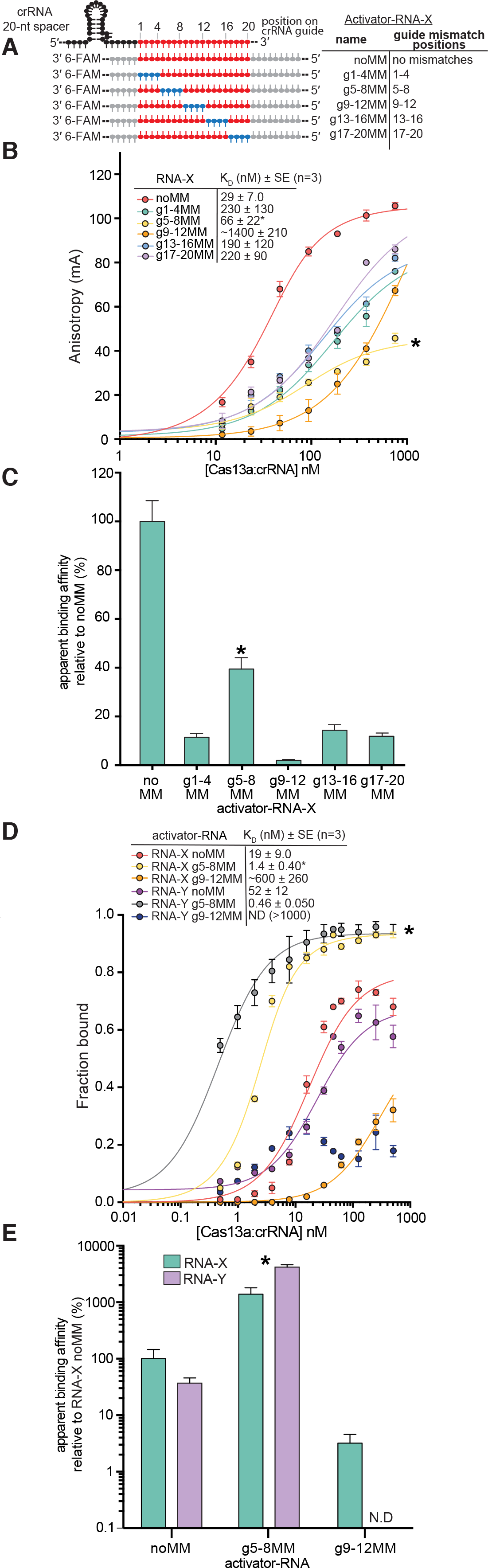
Biochemical validation of Lbu-dCas13 binding profiles. (**A**) Schematic of the Lbu-Cas13a crRNA: ssRNA activator interaction highlighting the different activator-RNAs tested in this study. 6-FAM: a covalently attached 6-carboxyfluorescein fluorophore. (**B**) Fluorescence anisotropy binding curves for the interaction between Lbu-dCas13a: crRNA-X and X activator-RNAs depicted in (A). Binding data fits (see Methods) shown as solid lines. Error-bars represent the s.d of the anisotropy from three independent experiments. Dissociation constants (K_D_) and their associated standard errors (SE) from three independent experiments are shown. (**C**) Data from (B) normalized as a percentage binding affinity relative to the affinity to a perfectly complementary activator-RNAs (no MM). (**D**) filter binding curves for the interaction between Lbu-dCas13a: crRNA-X (or Y) and X (or Y) 5’ ^32^P-labelled and 3’-6FAM fluorophore-labelled activator-RNAs. ^32^P phosphor-imaging was used to detect the activator-RNAs in this experiment. Binding data fits (see Methods) shown as solid lines. A binding curve could not be fit RNA-Y g9-12MM. Error-bars represent the s.d of the fraction bound from three independent experiments. Dissociation constants (K_D_) and their associated standard errors (SE) from three independent experiments are shown in the Fig. (**E**) Data from (D) normalized as a percentage binding affinity relative to the affinity to a perfectly complementary activator-RNA X (no MM) with error bars representing the normalizeds.d from three independent experiments. In all panels, g5-8MM binding data is marked with an (*).

Because our sequencing assay hinted that pairs mismatches at positions 5-8 (at for RNA-X) could somehow compensate for the binding defect seen for the analogous singly-mismatched sequences, we were interested in understanding the effect of four contiguous mismatches at positions 5-8 (g5-8MM). To our surprise, we noticed that in comparison to other sets of mismatches, g5-8MM exhibited far more moderate ~two-fold reduction in binding affinity relative to compared noMM target, in our fluorescence anisotropy assay **(Fig. 3B, C)**. Peculiarly, we also noticed that the overall change in fluorescence anisotropy signal between unbound and fully bound activator-RNA was substantially lower (at least two-fold) for g5-8MM compared to noMM activator-RNAs **(Fig. 3B**, see binding curve marked with an asterisk). This atypical effect was reproducible across three different RNA sequences **(Fig S4A)**. Thus, we speculated that the g5-8MM result might be artefactual, possibly a result of the 3’ fluorophore exhibiting significantly more rotational flexibility upon binding leading to an inaccurate estimation of binding affinity.

In order to disentangle this potentially confounding effect from the binding behavior of targets with mismatches at positions 5-8, we performed (fluorescence-independent) filter-binding experiments with Lbu-dCas13a: crRNA complexes and ^32^P 5’-radiolabelled activator-RNAs. In order to be as consistent as possible with the fluorescence anisotropy data, the targets used in these experiments also contained the 3’ fluorophore chemical modification. Using this setup, we observed good concordance of binding affinities measured by fluorescence anisotropy and filter binding for Lbu-dCas13a: crRNA for noMM and g9-12MM activator (**Fig. 3D, E)**. These results were consistent across two different crRNA: target RNA sequences (RNA-X and RNA-Y; **Fig. 3D, E).** However, and to our surprise, we measured a binding affinity approximately ~10-100-fold tighter for g5-8MM compared to noMM activator-RNAs (**Fig. 3D-E**, see data marked with an asterisk). To further verify that the inconsistency between observed binding affinities for g5-8MM targets was due to an artifact in the fluorescence anisotropy experiment, we used radiolabeled filter binding experiments to measure binding affinities for targets with or without at 3’ fluorophore modification. The observed binding affinities were not affected by the 3’ fluorescent label across three different RNA sequences (RNA-X, Y and Z.) **(Fig. S4B)**.

Taken together these experiments validate the compensatory effects of single mismatches between nucleotides 5-8 that we hypothesized from the sequencing data, and that a block of mismatches at nucleotides 5-8 can appear to promote even tighter Cas13a-RNA binding relative to a perfectly complementary crRNA-guided Cas13a: activator-RNA interaction across a range of RNA sequences.

### Cas13 binding profiles are unaffected by the length of the crRNA: target duplex

Although we performed our high-throughput sequencing assay on a fixed guide length of 20nt, we were also interested in understanding the extent to which the observed mismatch-tolerance profile was affected by varying the length of the crRNA guide. This was motived by the observation that that Cas13a’s HEPN-nuclease activity can be activated using a range crRNA guide sequence lengths (Abudayyeh et al., 2017; Abudayyeh et al., 2016; Cox et al., 2017; East-Seletsky et al., 2016; East-Seletsky et al., 2017; Gootenberg et al., 2017; Knott et al., 2017; Konermann et al., 2018; Liu et al., 2017a; Smargon et al., 2017; Yan et al., 2018), and led us to perform the same fluorescence-anisotropy binding assay with an analogous set of fluorophore-labeled mismatched activator-RNAs and a crRNA containing a 24-nt. guide sequence **(Fig. S4C-D).** We found that extending the crRNA: activator-RNA duplex to 24-nt did not increase the overall affinity of the Cas13a: activator-RNA interaction compared to a 20-nt crRNA **(Fig. S4D-E).** This observation suggests that either the additional base-pairing interactions do not occur or are energetically neutral with respect to binding affinity between a crRNA and its perfectly complementary activator-RNA. Furthermore, the consistency of mismatch (in)tolerances between 20- and 24-nt crRNA duplexes indicate that the results we observe through the high-throughput sequencing experiments are not an artifact of the specific guide length used for the experiment, but rather a general property of Lbu-dCas13a. Taken together, these data add to the evidence that LbuCas13a’s binding affinity to activator-RNA is determined by a number of unequal and complex contributions from individual base-pairing interactions.

### Mismatched target RNAs differentially impact Cas13a HEPN-nuclease activation

Next, we wondered whether mismatched activator-RNAs might still activate Cas13a for trans-RNA cleavage directed by the HEPN-nuclease, and particularly whether we would observe mismatch position-dependent effects. To measure Cas13a HEPN-nuclease activation, we used a fluorescent RNA cleavage assay (East-Seletsky et al., 2016) that assesses HEPN-mediated RNA cleavage upon RNA-mediated Cas13 activation (**Fig. 4A**). The multiple turnover conditions of this assay coupled with the robust catalytic turnover of Lbu-Cas13a allows us to detect HEPN-nuclease activity from as little as ~10pM of activator-RNA bound Cas13a: crRNA complex (East-Seletsky et al., 2016), enabling the rapid measurement of a range of parameters that alter the HEPN-nuclease activity of Cas13a.

**Fig. 4.**
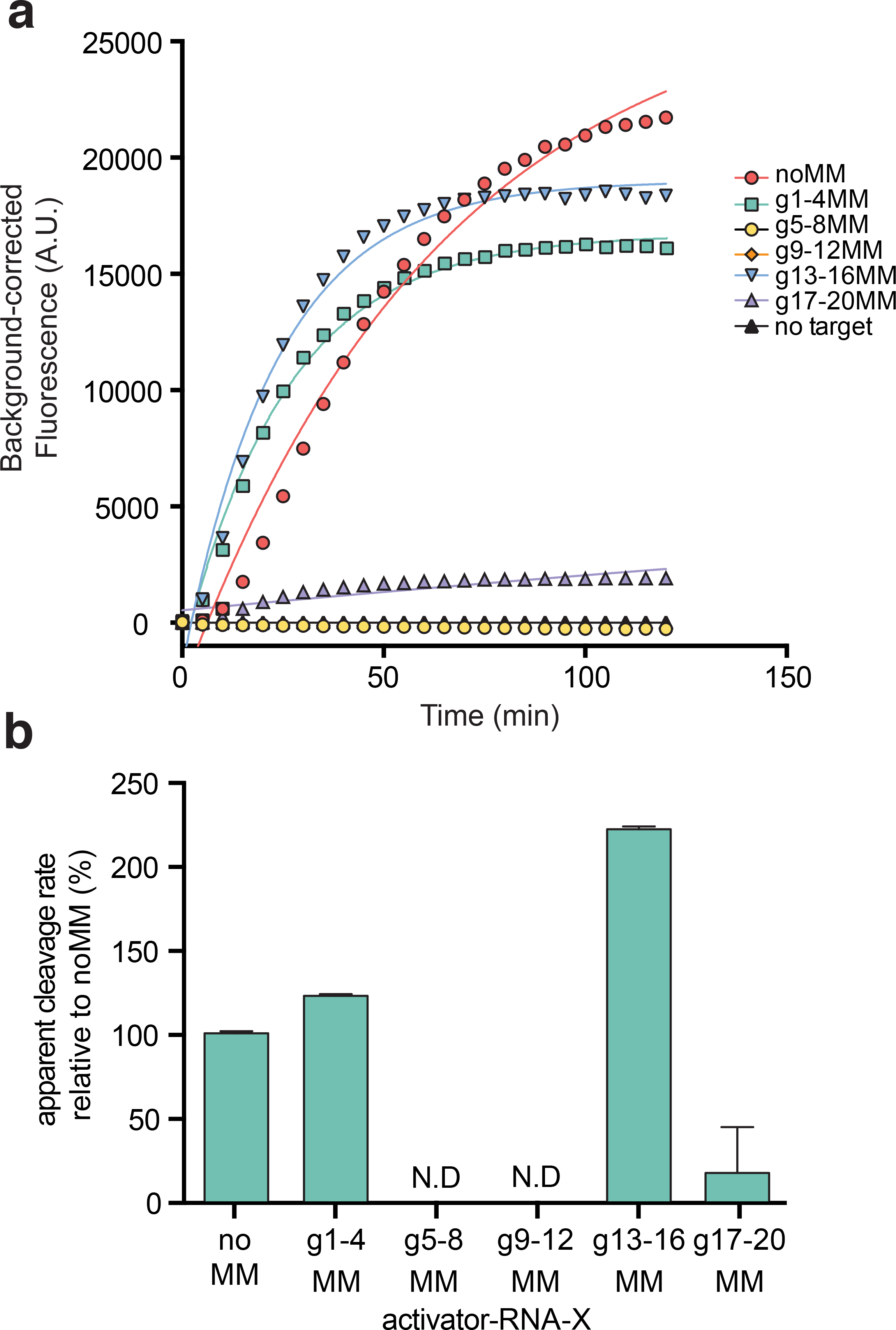
Position of crRNA: activator-RNA mismatches differentially impact LbudCas13a HEPN-nuclease activation. (**A**) representative time course of background corrected fluorescence measurements generated by Lbu-Cas13a: crRNA-X activation by the addition of 100 pM X activator-RNAs with either zero (noMM) or four consecutive mismatches across the crRNA-spacer targeting region. Quantified data were fitted with single-exponential decays (solid line) with calculated pseudo-first-order rates constants (*k*_obs_) (mean ± s.d., n =3) as follows: noMM 0.023 ± 0.003 min^-^1, g1-4MM 0.028 ± 0.002 min^-^1, g13-16MM 0.050 ± 0.003 min^-^1, g17-20MM 0.004 ± 0.007 min^-^1, while g5-8MM and g9-12 were not fit. (**B**) Data from (A) normalized as a percent cleavage rate relative to a perfectly complementary activator-RNA X (no MM) with error bars representing the normalized s.d from three independent experiments.

Using this assay, we determined relative apparent rates of fluorescent RNA reporter cleavage (normalized to the completely complementary RNA-activator; noMM) from each of the resulting time-courses (**Fig. 4B**). The data revealed a range in the ability of each mismatched activator-RNA to activate Cas13a trans-RNA cleavage, and similar effects were observed using a crRNA with a 24-nt guide sequence and a corresponding set of activator-RNAs (**Fig. S5A-B)**.

Despite exhibiting weaker binding to the Cas13a: crRNA complex, RNAs g13-16MM and g1-4MM are still able to promote robust Cas13a HEPN-nuclease activity, with cleavage rates greater or similar to the catalytic rates promoted by a perfectly complementary activator-RNA (noMM), respectively (**Fig. 4B**). This observation suggests that under the assumed equilibrium conditions of this assay, any decrease in the amount of active activator-RNA-bound Cas13a complex (due to a reduced affinity) is offset by increased catalytic turnover of the HEPN-nuclease when mismatches are present in these regions. On the other hand, some quadruple-mismatch activator-RNAs produced large reductions in Cas13a HEPN-nuclease activation (RNAs g5-8MM, g9-12MM & g17-20MM). The defects in Cas13a catalytic activation exhibited by RNAs g9-12MM and g17-20MM could be either attributed to their weak binding affinities for Cas13a: crRNA (and the lack of a sufficient amount of active activator-RNA-bound Cas13a complex to create a fluorescent signal) or a specific requirement for a base-paired RNA duplex in these regions for HEPN-nuclease activation.

Most surprisingly, we found that activator-RNAs with mismatches at guide positions 5-8 (g5-8MM) are unable to trigger the Cas13a HEPN-nuclease activity (**Fig. 4B**), even though they are bound more tightly by Cas13a: crRNA than a perfectly-complementary activator-RNA (**Fig. 3**). We confirmed that this phenomenon occurs across a number of different crRNA: activator-RNA sequences, and that HEPN-nuclease activation occurs upon restoration of base-pairing interactions between the crRNA and the activator-RNA in this region (**Fig. S5C-F)**. This striking result indicates that stable activator-RNA binding and HEPN-nuclease activation can be decoupled. Furthermore, the conformational rearrangement of the HEPN-domains required to form a catalytically competent nuclease active site requires a base-paired RNA duplex at positions 5-8 of the crRNA spacer. Together, these results reveal a complex interplay between crRNA-spacer: activator-RNA base-pairing requirements for stable Cas13a binding and for HEPN-nuclease activation.

### RNA base-pairing is an allosteric activator of Cas13a HEPN-nuclease activity

To determine how crRNA: activator-RNA base pairing triggers Cas13a HEPN-nuclease catalytic activity, we tested whether activator-RNAs containing single nucleotide mismatches to the crRNA guide sequence can activate Cas13a for non-specific RNA cleavage (**Fig. 5A & Fig. S6)**. Overall, Cas13a nuclease activation was more affected by single-mismatches in the guide-RNA nucleotide positions 5-8 and 17-20 compared to the positions 1-4, 9-12 and 13-16 (**Fig. 5A**). The largest defect in activation was caused by a mismatch in guide-RNA position 6, which resulted in a fivefold reduction in cleavage rate compared to a perfectly complementary target RNA. This indicates that a single base-pair mismatch within the crRNA: target RNA duplex is sufficient to impair Cas13a HEPN-nuclease activation, despite having complex effects on Cas13a binding affinity (**Figs. 2-3**).

**Fig. 5.**
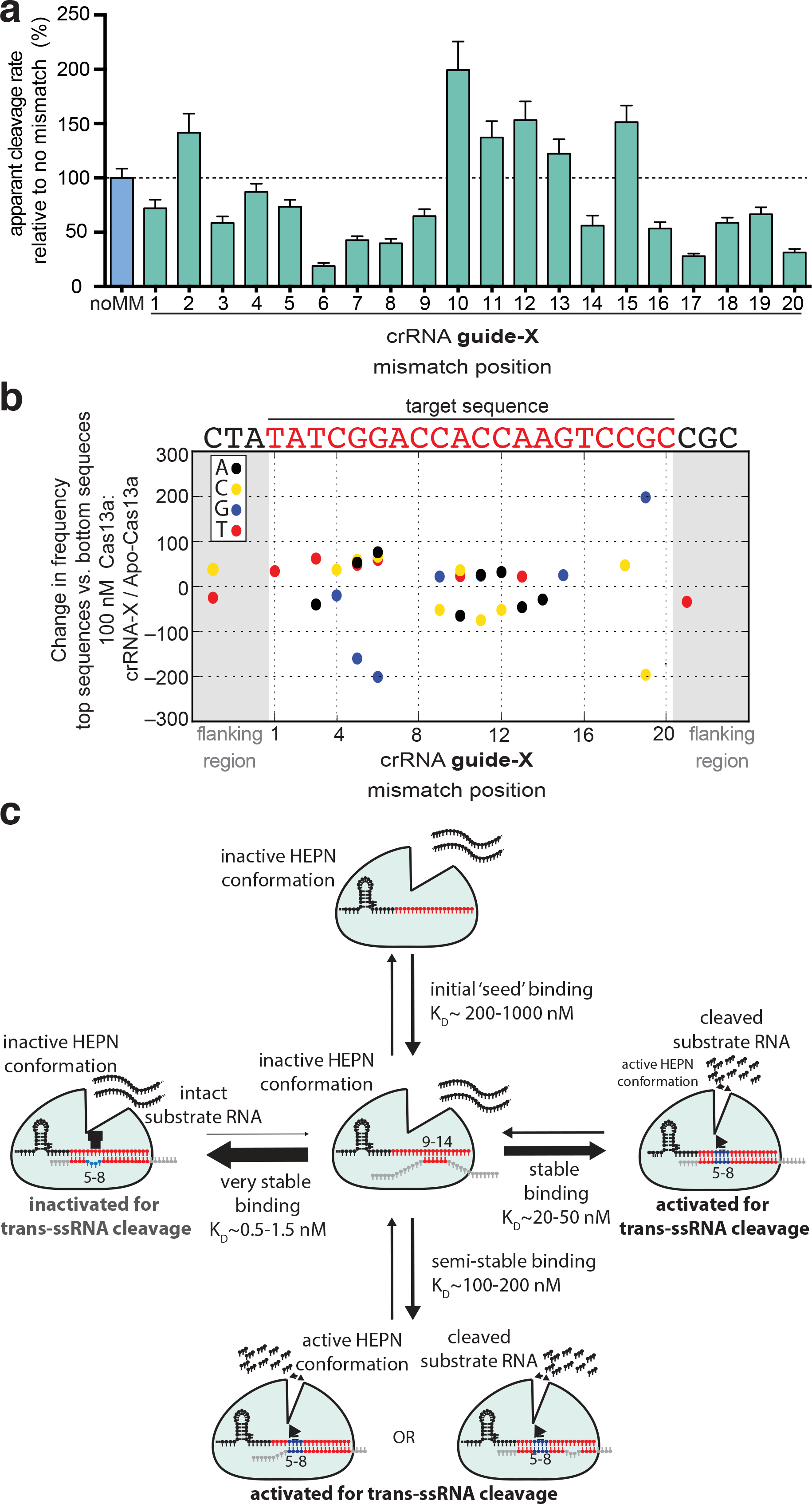
Lbu-Cas13a HEPN-nuclease activation is gated by ‘unfavorable’ base-pairing within the context of the Cas13a crRNA: activator-RNA interaction. Background corrected apparent cleavage rates generated by Lbu-Cas13a: crRNA-X activation by the addition of 100 pM activator-RNA X with either zero (noMM) or single mismatches in positions 1-20 across the crRNA-spacer. Apparent cleavage rates are plotted as a normalized percentage cleavage rate relative to a perfectly complementary activator-RNA X (noMM) with error bars representing the normalized s.d from three independent experiments. Representative time-course data can be found in Supplementary Data Figure 9A. (**B**) Change in nucleotide frequency in each position shown between the top 500 enriched members and the bottom 500 enriched members (as determined by fold-change relative to Apo Lbu-dCas13a) for 100 nM dCas13a: crRNA-X targeting the activator-RNA X library. The activator-RNA X crRNA-target sequence is shown above. Values with p ≥ 0.001 in the permutation test are not shown. (**C**) A revised model for Lbu-Cas13a HEPN-nuclease activation based on the key findings in this study.

Conversely, three out of four mismatches within the 9-12 region resulted in more robust Cas13a HEPN-nuclease activation relative to perfectly complementary RNA target sequences, indicating that single mismatches in this region do not preclude HEPN-activation. In fact, these target RNAs promote higher catalytic activity (**Fig. 5A**). Comparing this result to the high-throughput mismatch profiling data (**Fig. 2**) where mismatches in this region had the greatest negative effect on binding affinity, reiterates that base-pair mismatches in some cases may act to improve HEPN-nuclease catalytic activity, despite their negative effect on binding affinity.

Together, these two observations prompted us to revisit the high-throughput mismatch profiling data to see whether implementing a ranking-based approach would detect enrichment or depletion of base-pairing interactions regardless of whether the crRNA: target RNA duplex contained single or multiple mismatches. Specifically, we ranked all RNA-X library members from the 100 nM Cas13a: crRNA-X sample based on their fold-change enrichment / depletion compared to the apo-Cas13a sample. We then separately computed the nucleotide frequency by position for the top 500 members and the bottom 500 members of the list and used this to compute the frequency difference between the two subsets (**Fig. 5B**). To deal with any possible heterogeneity in the background nucleotide distribution, we performed permutation tests using shuffled lists of sequences as a null hypothesis (see **Methods**). This analysis identified several positions within the crRNA: activator-RNA duplex that exhibit interesting nucleotide preferences. Most striking was the substantial depletion at positions 5 and 6 of guanine nucleotides that are complementary to the crRNA, and a slight enrichment of the remaining three nucleotide identities at these positions. This observation indicates that Cas13a: crRNA binds more tightly on average, to members of the RNA library that allow for mismatches in g5 and g6 crRNA positions and the target RNA, even in the presence of other mismatches, in agreement with the filter-binding data in Fig. 3 again suggesting the presence of non-additive, compensatory binding effects when multiple guide: target mismatches are present.

## DISCUSSION

Our investigation of specificity requirements for CRISPR-Cas13a RNA-activated RNA cleavage found that mismatches between crRNA and activator-RNA affect RNA binding affinity and HEPN-nuclease activation differently depending on mismatch position. These results reveal a mechanism following initial crRNA: activator-RNA base pairing that serves as a specificity checkpoint controlling HEPN-nuclease activation (**Fig. 5C).** Our data support a model for Cas13a target search that involves activator-RNA base-pairing with a ‘mismatch hyper-sensitive’ or ‘seed’ region (guide-RNA nucleotides 9-14) and a ‘HEPN-nuclease switch region (guide-RNA nucleotides 5-8). Target search by crRNA-loaded LbuCas13a most probably requires activator-RNA base-pairing in the mismatch hyper-sensitive region, and stable binding requires additional contributions from supplementary base-pairing at guide-RNA positions 1-4 and 15-20. After the activator is bound, HEPN-nuclease activation is gated in part by LbuCas13a detecting base-pairing between the crRNA-guide and the HEPN-nuclease switch region of the activator-RNA, triggering HEPN-domain conformational rearrangement and formation of a catalytically competent active-site (Liu et al., 2017a; Liu et al., 2017b). Mismatches within the HEPN-nuclease switch region may block conformational change by trapping Cas13a in a non-catalytic yet more tightly-bound activator-RNA-associated state. We propose that multiple mismatches between activator-RNA nucleotides and guide RNA at positions 5-8 bypass an energetically unfavorable rearrangement that occurs when a fully complementary RNA activator binds Cas13a. Conversely, it appears that even though multiple mismatches in guide RNA positions 1-4 and 13-16 are energetically unfavorable with respect to binding affinity, they are still able to promote robust HEPN-nuclease activity. This model is supported by data for both Lsh-Cas13a (Abudayyeh et al., 2016) and Lwa-Cas13 (Gootenberg et al., 2017) where multiple mismatches in guide RNA regions 5-8 and 9-12 compromise HEPN-nuclease activity. Furthermore, our data and model rationalize the mismatch scheme developed to maximize SNP detection sensitivity within the Lwa-Cas13a RNA detection platform (Gootenberg et al., 2017). Taken together, these similarities suggest broad conservation of activator-RNA binding and HEPN activation mechanisms (and their sensitivity to mismatches) across the U-cleaving Cas13a family. Subtle variation in the magnitude or position of these effects may exist between homologs, given the broad range of activator sensitivities and rates of *trans*-ssRNA cleavage observed across this family (Abudayyeh et al., 2016; East-Seletsky et al., 2016; East-Seletsky et al., 2017; Gootenberg et al., 2018; Gootenberg et al., 2017).

A number of nucleic-acid guided proteins employ accessible, pre-ordered ‘seed’ regions within their guides for target search (reviewed in (Gorski et al., 2017; Jackson et al., 2017)). The binary structures of Lsh-Cas13a (Liu et al., 2017b), Lbu-Cas13a (Liu et al., 2017a), and Lba-Cas13a (Knott et al., 2017), show the repeat proximal spacer nucleotides adopting a distorted ‘U-turn’ conformation within the nuclease lobe, while the repeat distal spacer nucleotides are solvent-exposed. Examining the crRNA-loaded structure of Lbu-Cas13a (Liu et al., 2017a) shows that the ‘mismatch sensitive’ region we identify in this study is solvent-exposed in a near A-form conformation, but engaged in mismatched base-pairing with the 3’-end of the spacer. While the apparent pre-ordering and artefactual mismatch binding suggests this could be a seed for activator RNA binding, our data definitively identifies these nucleotides as the seed for Lbu-Cas13a.

The structure of Lbu-Cas13a: crRNA bound to activator-RNA (Liu et al., 2017a) combined with our mismatch profiling, together provide the basis to rationalize how the HEPN-nuclease switch region might allosterically regulate the HEPN-nuclease. Both the 3’ flank of the crRNA-repeat and the HEPN-1 domain undergo a concerted conformational change that drives the formation of a catalytically competent active-site (Liu et al., 2017a). Of note is an alpha helix of the HEPN-1 domain that is closely packed across the phosphate backbone of the HEPN-nuclease switch region activator RNA nucleotides 5*-8* (complementary to g5-8; **Fig. 6**). In particular, S786 interacts through a bidentate hydrogen bonding interaction with a non-bridging oxygen from the phosphate of activator-RNA nucleotide 6* and C(-1) within the 3’ flank of the crRNA-repeat. This interdependent sensing of activator-RNA nucleotide 6* by the crRNA 3’ flank C(-1) via the HEPN-1 domain may drive conformational activation of the HEPN-nuclease. Consistent with this, mismatches in this region would shift the positions of the activator RNA phosphates, disrupting the geometry of the crRNA: activator duplex leading to conformational inhibition of the HEPN-nuclease. This network of interactions might have implications for the mechanism by which the PFS (Abudayyeh et al., 2016) is able to gate HEPN-nuclease activity, given that C(-1) has been implicated in sensing the PFS (Liu et al., 2017a; Liu et al., 2017b). In addition to this interaction, a non-bridging oxygen from the phosphate of crRNA nucleotide 7, which base-pairs with nucleotide 7* in our activator-RNA, interacts with K558 from the Helical-2 domain, and mutation of this residue to alanine resulted in a significant defect in HEPN-nuclease activity (Liu et al., 2017a). Based on our data, one can imagine that the role of K558 and S786 is to simultaneously sense the positions of crRNA and activator-RNA phosphates, respectively, relative to one another only when they exist in an RNA duplex. It is worth noting that even single mismatches in an RNA:RNA duplex can have large effects on the local geometry of the duplex including positions of the phosphate groups (Pan et al., 1998). Further studies will be required to understand how the dynamics of Cas13a: crRNA complexes allow them to overcome the energetic barriers imposed by crRNA organization to drive activator-RNA binding, and how sensing of the RNA duplex in this region by the helical-II and HEPN-1 domains, as well as the crRNA is able to elicit HEPN-nuclease activation.

**Fig. 6.**
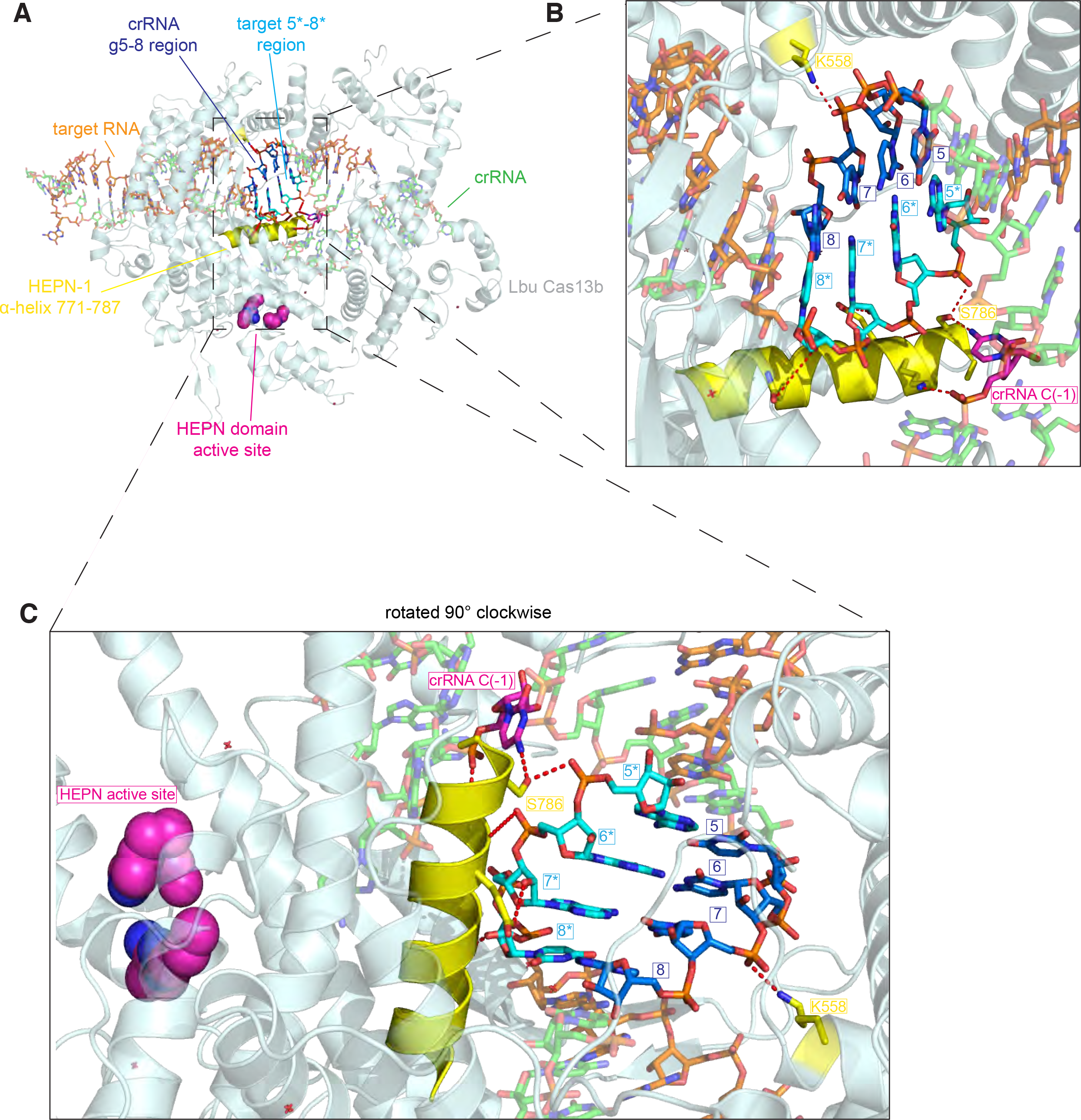
Putative structural basis of Cas13a detection of crRNA: target-RNA base-pairing in the g5-8 region. Overall view of the ternary Cas13a: crRNA:target-RNA crystal structure. Features implicated in the recognition of the crRNA-spacer nucleotides 5-8: target nucleotide 5*-8* (complementary to g5-8) RNA duplex region are labeled. (**B)** Close-up view of the crRNA g5-8: target 5*-8* (complementary to g5-8) RNA duplex region with important features labeled. (**C**) same as B, except the ternary complex has been rotated 90° clockwise. Figure generated from PDB ID:5XWP.

Despite a fundamental difference in guide RNA organization, there are some striking similarities between Cas13a and Argonaute (Ago) proteins with respect to target ssRNA recognition and catalytic activation. First, relative to perfectly complementary targets, mismatches in guide nucleotides 9-12 improve mouse Ago2’s affinity for target ssRNA (Wee et al., 2012) much like what we observed with mismatches in crRNA region 5-8 of Lbu-Cas13a. Second, base-pairing in this region is required for Ago2 proteins to achieve a catalytically competent ‘slicer’ conformation active for nucleic acid cleavage (Haley and Zamore, 2004; Martinez and Tuschl, 2004; Schwarz et al., 2006), similar to HEPN-nuclease activation within Cas13a. And finally, while Ago2 proteins in animals have retained their ability to cleave perfectly-complementary target RNAs (i.e. through the siRNA pathway), they primarily use mismatched guides (as describe above) to bind target RNA and recruit host factors to carry out translational repression of target mRNAs(i.e. the miRNA pathway) (Bartel, 2009). It is tantalizing to speculate that this function could be used to modulate the expression of viral or self mRNAs upon viral infection, and/or during normal function, or modulate to virulence, much like the CRISPR-Cas9 system from *Francisella novicida* (Sampson et al., 2014; Sampson et al., 2013). Alternatively, similar to other CRISPR-Cas systems, there might be a role for mismatched crRNA-spacer: ssRNA targets in the acquisition phase of CRISPR immunity, akin to primed spacer acquisition in type I CRISPR systems (Datsenko et al., 2012; Fineran et al., 2014). Additionally, our observation that mismatches in some cases seemed to be tolerated, if not preferred with respect to maximal HEPN-nuclease activity, which might be indicative of an evolved mismatch tolerance required to maintain immunity when faced with the rapidly evolving nature of phage genomes. Indeed, recent data suggests that RNA-targeting by type III CRISPR-Cas systems exhibit relaxed interference specificity relative to type I and II DNA targeting CRISPR-Cas systems (Pyenson et al., 2017). Nevertheless, further work is required to understand whether there is a role for both of these behaviors within the native hosts of type VI CRISPR-Cas systems.

Understanding the specificity of Cas13a activator-RNA binding and subsequent HEPN-nuclease activation is crucial for all current (Abudayyeh et al., 2017; Aman et al., 2018; Cox et al., 2017; East-Seletsky et al., 2016; East-Seletsky et al., 2017; Gootenberg et al., 2018; Gootenberg et al., 2017; Konermann et al., 2018) and potential applications of Cas13a’s programmable RNA-binding and/or cleavage capabilities. In particular, recent reports of specific RNA-cleavage in plant (Abudayyeh et al., 2017; Aman et al., 2018) and human (Abudayyeh et al., 2017; Cox et al., 2017; Konermann et al., 2018) cell lines, as well as RNA-editing (Cox et al., 2017) and modulation of premRNA splicing (Konermann et al., 2018) in human cell lines underscore the need to understand how Cas13 is able to specifically bind and in some cases cleave specific RNA targets while avoiding similar sequences, across a range of organisms. Our data enable rational design of crRNAs with optimal specificity and activity for each type of application, and underscore a need to consider the range of effects that Cas13a off-target recognition might have on Cas13a’s ssRNA-binding and/or cleavage behavior, and ultimately the fate of the RNA pool present in each experiment.

## METHODS

### Lbu-dCas13a-Avitag overexpression and purification

HEPN-nuclease deactivated Lbu-Cas13a (dCas13a: R472A, H477A, R1048A, H1053A; Addgene plasmid #: 83485) was subcloned into pDuBir (a generous gift from Laura Lavery) in order to express a Lbu-dCas13a with a *N-*terminal His_6_-MBP-TEV cleavage site tag, a C-terminal BirA biotinylation signal (Avitag(Beckett et al., 1999)) as well as a separate BirA polypeptide, which biotinylates the Avitag. Expression of both polypeptides is driven by a T7 promoter. pDuBir-Lbu-dCas13a-avitag (Addgene plasmid #: 100817) was transformed into Rosetta2 *E. coli* cells grown in Terrific broth at 37 °C. *E. coli* cells were induced at OD_600_ ~1 with 0.5 M ITPG and 50 µM D-biotin (in 10 mM bicine buffer pH8.3) was added as substrate for the biotinylation reaction. The temperature was then reduced to 16 °C for overnight expression of His-MBP-Cas13a-Avitag. Cells were subsequently harvested, resuspended in lysis buffer (50 mM Tris-HCl pH 7.0, 500 mM NaCl, 5% glycerol, 1 mM TCEP, 0.5mM PMSF, 50 µM D-biotin, and EDTA-free protease inhibitor (Roche)), lysed by sonication, and the lysates were clarified by centrifugation. Soluble His-MBP-Cas13a-Avitag was isolated by Ni-NTA affinity chromatography. While concurrently dialyzing into ion exchange buffer (50 mM Tris-HCl pH 7.0, 250 mM KCl, 5% glycerol, 1 mM TCEP), the His-MBP-dCas13a-Avitag containing eluate was incubated with TEV protease at 4 °C overnight in order to cleave off the His_6_-MBP tag. dCas13a-Avitag was loaded onto a HiTrap SP column and eluted over a linear 0.25-1M KCl gradient. Cation exchange chromatography fractions containing dCas13a-Avitag were pooled and further purified via size-exclusion chromatography on an Superdex S200 column and stored in gel filtration buffer (20 mM Tris-HCl pH 7.0, 200 mM KCl, 5% glycerol, 1 mM TCEP) at -80C until required. Biotinylation efficiency was determined using a streptavidin-magnetic bead pulldown experiment, as described in ref. (O’Connell et al., 2014). Wildtype Lbu-Cas13a (Addgene plasmid #: #83482) was expressed and purified as previously described (East-Seletsky et al., 2016), essentially as above except D-biotin wasn’t added during overexpression and purification.

### Preparation of RNA

All 3’-prime fluorescein (6-FAM) labelled activator-RNAs were purchased from Integrated DNA technologies (IDT). All crRNAs and unlabeled activator-RNAs were transcribed *in vitro* from either PCR-amplified double-stranded DNA templates (crRNAs) or single-stranded DNA templates with an annealed dsDNA T7 promoter region (IDT). All templates contained a sequence encoding a hepatitis delta virus (HDV) ribozyme 3¢ to the desired RNA product to ensure 3’ end homogeneity. Transcriptions were carried out as described previously(Sternberg et al., 2012) with the following modifications: RNAs were transcribed at 37°C for 3 h, followed by treatment with RQ1 RNase-free DNase (Promega) and 5 mM MgCl_2_ (37°C, 1 hr) to remove DNA templates and drive ribozyme cleavage, respectively. Transcription reactions were purified by 15% denaturing polyacrylamide gel electrophoresis (PAGE), activator-RNAs were resuspended in cleavage buffer (20 mM HEPES pH 6.8, 50 mM KCl, 5 mM MgCl_2_, and 5% glycerol), while crRNAs were first resuspended in RNase-free water, then prior complexing with Lba-Cas13a, crRNAs were folded in 1X annealing buffer (10 mM Na-HEPES pH 7.4, 30 mM KCl, 1.5 mM MgCl_2_) by heating at 75°C for 5 min followed by rapid cooling on ice. For filter-binding experiments, 5’ triphosphates were removed from activator-RNAs by calf intestinal phosphate (New England Biolabs) prior to radiolabeling and activator-RNAs were then 5’-end labeled using T4 polynucleotide kinase (New England Biolabs) and [γ-^32^P]-ATP (Perkin Elmer) as described previously(Sternberg et al., 2012). All oligonucleotide sequences used in this study can be found in Supplementary Data Table 2.

### Fluorescence anisotropy binding assays

Lbu-dCas13a-avitag:crRNA complexes were preassembled by incubating 1μM of Lbu-dCas13a with 1.1 µM of crRNA in modified RNA processing buffer (20 mM HEPES pH 6.8, 50 mM KCl, 5 mM MgCl_2_, 10 μg ml^−^1 BSA, 100 μg ml^−^1 yeast tRNA, 0.01% Igepal CA-630 and 5% glycerol) for 2 min at 37°C, followed by 8 min at 25°C. These complexes were then serially diluted multiple times in modified RNA processing buffer to evenly cover between 0 - 1 µM Lbu-dCas13a: crRNA. These complexes were then further diluted two-fold in modified RNA processing buffer in the presence of 40 nM (final concentration) of 3’-labelled 6-FAM activator-RNA. These reactions were incubated for one hour at 37°C before fluorescence anisotropy measurements were taken (λ_ex_: 485nm; λ_em_: 535 nm). Fluorescence anisotropy values (mA) were fit using a solution of the quadratic binding equation using GraphPad Prism, to avoid any effects of protein depletion when [RNA] > K_D_ (see equation (6) in ref. (Roehrl et al., 2004)). For each curve fit, the maximum anisotropy value was floated to allow for activator-RNA specific effects on magnitude of maximum anisotropy value. All experiments were carried out in triplicate. Dissociation constants and associated errors are reported in the Fig.s and/or Fig. legends.

### Filter-binding assays

Lbu-dCas13a-avitag:crRNA complexes were preassembled as described above. These complexes were then serially diluted multiple times in modified RNA processing buffer to evenly cover between 0 - 1 µM Lbu-dCas13a: crRNA. These complexes were then further diluted two-fold in modified RNA processing buffer in the presence of ~0.5 nM (final concentration) of 5’-^32^P-labelled activator-RNA. These reactions were incubated for one hour at 37°C prior to loading samples onto a dot-blot apparatus. Tuffryn, Protran and Hybond-N+ were assembled onto a dot-blot apparatus in the order listed above. Prior to sample application, each membrane was washed twice with 50μL Equilibration Buffer (20 mM HEPES pH 6.8, 50 mM KCl, 5 mM MgCl_2_ and 5% glycerol). 25 µL of each sample was loaded per well, followed by a single 50 μL Equilibration Buffer wash. Membranes were then briefly dried and visualized by phosphorimaging (Typhoon FLA9500 scanner, G.E Healthcare). Fraction-bound values were fit using a solution of the quadratic binding equation using GraphPad Prism, to avoid any effects of protein depletion when [RNA] > K_D_ (see equation (6) in ref. (Roehrl et al., 2004)). For each curve fit, the max fraction bound value was constrained be greater than 0.6 to ensure (a) that weakly bound activator-RNAs affinities where not overestimated, and (b) affinities were not underestimated for activator-RNA interactions where the fraction bound values plateaued at less than 1.0. All experiments were carried out in triplicate. Dissociation constants and associated errors are reported in the Fig.s and/or Fig. legends.

### Fluorescent ssRNA reporter nuclease assays

LbuC2c2:crRNA complexes were preassembled by incubating 1μM of WT Lbu-Cas13a with 500 nM crRNA in modified RNA processing buffer for 2 min at 37°C, followed by 8 min at 25°C. These complexes were then diluted to 100nM LbuC2c2: 50 nM crRNA in modified RNA processing buffer in the presence of 150 nM of RNAase-Alert substrate (Thermo-Fisher), 100 ng of HeLa total RNA and 100 pM of activator-RNA. These reactions were incubated in a fluorescence plate reader for up to 120 min at 37°C with fluorescence measurements taken every 5 min (λ_ex_: 485 nm; λ_em_: 535 nm). Background-corrected fluorescence values were obtained by subtracting fluorescence values obtained from reactions carried out in the absence of target ssRNA. Background normalized apparent cleavage rates were calculated using a single-exponential decay using GraphPad Prism and associated errors are reported in each Fig. as standard deviation error bars from three independent experiments.

### High-throughput mismatch profiling library design rationale and preparation

Two distinct ssDNA 100-nt libraries containing a 34-nt ‘target-region’ comprising a partially-randomized library (76% frequency of on-target nt. and 8% frequency of each other nt. in each position) of either RNA X or Y sequences, flanked by Illumina sequencing adaptor complementary sequences, a random 6-nt. ‘Cluster ID’ region for efficient cluster calling, and a 5’ T7 promoter for *in vitro* transcription (**Fig. 3A**). The 34-nt target should theoretically cover all single- and pairwise-error combinations, as well as a substantial portion of triple-mismatch combinations (see Simulate_fragments.ipynb script github.com/OConnell-lab-URMC/High-throughput-RNA-mismatch-profiling to simulate mismatch distributions). Each library is barcoded twice: (1) during final PCR, a ‘P7’ index-containing adaptor was appended to allow for sample multiplexing within each sequencing experiment/run, and (2) during reverse transcription, a modified ‘P5’ adaptor containing a randomized 13-n.t. ‘unique molecule identifier’ was appended to correct for any library PCR bias. Each library was produced by in vitro transcription as described above from single-stranded DNA template (IDT) annealed to a short ssDNA to create a dsDNA T7 promoter region.

### High-throughput mismatch profiling ssRNA library dCas13a-avitag pull-down

Lbu-dCas13a-avitag:crRNA complexes were preassembled as described above (excepted for Apo-Lbu-dCas13a-avitag samples, where the crRNA addition was omitted). X and Y activator-RNA libraries (final concentration 100 nM of each) were added to modified RNA processing buffer with 40 U RNasin (Promega), and Lbu-dCas13a-avitag:crRNA complexes (10 or 100 nM Lbu-dCas13a-avitag) in a total volume of 100 µL and incubated at 37 °C for 1 h. This mixture was then added to 12 µL magnetic streptavidin beads (Dynabeads MyOne Streptavidin C1; Life Technologies) pre-equilibrated in modified RNA processing buffer and agitated at 4 °C for 1 h. Beads were then washed once with 1 mL wash buffer (20 mM Tris-HCl, pH 7.5, 50 mM NaCl, 5mM MgCl_2,_ 0.1% Triton X-100, 5% glycerol, 1mM DTT). Bound RNAs were eluted from dCas13a-avitag by DEPC-water/trizol extraction, and purified by phenol-chloroform extraction and ethanol precipitation.

### High-throughput mismatch profiling Illumina sequencing library construction

RNA pellets from the library pulldown were resuspended with 15µL RNase-free water. 33% of the sample (or 0.5 pmol X and Y activator-RNA library as input control) was annealed with 2pmol RT primer (see Supplementary Data Table 2 for sequence) and reverse transcribed using Superscript III reverse transcriptase (Invitrogen) following the manufacturer’s protocol. The RT primer contains a 13-nt randomized sequence (instead of a standard P7 index) which serves as a unique molecular identifier (UMI) to enable accurate molecule counting. Excess primers were degraded with Exonuclease I (New England Biolabs), and products were purified by ethanol precipitation. cDNAs were subjected to limited PCR amplification (12 cycles) with Phusion polymerase (New England Biolabs), using the NEBNext library barcoding kit to append Illumina sequencing handles and sample barcode identifiers to the library fragments. The amplified library was then purified and size-selected using AmpureXP beads (Beckman / Agencourt) with a 1:1 ratio of beads to sample. Library quality was verified by Bioanalyzer prior to high-throughput Illumina sequencing.

### High-throughput mismatch profiling Illumina sequencing

High-throughput RNA mismatch profiling library samples were pooled and sequenced on an Illumina HiSeq 4000 using RapidRun configuration, with 50bp single-end reads, standard P7 index cycles, and custom P5 index cycles to sequence the complete 13-mer UMI. Sequencing was performed at the Vincent J. Coates Genomics Sequencing Laboratory at UC Berkeley. All sequencing reads are available through the NCBI SRA at BioProject ID: PRJNA399726.

### High-throughput mismatch profiling sequencing analysis/bioinformatics

Sequencing data was analyzed using custom python scripts. All scripts and analysis presented here is publically available at github.com/OConnell-lab-URMC/High-throughput-RNA-mismatch-profiling. Reads were de-duplicated by counting instances of the 13 nucleotide unique molecular identifier, and de-duplicated reads were then assigned to one of the two (X and Y activator-RNA) target sequences based on Hamming distance. The number of unique observations of each sequence variant was computed, and entries with low counts (fewer than 20 unique counts in each of 3 replicates for any given condition) were filtered out. Counts were normalized to the total number of unique reads in a given sample. Fold change between conditions was calculated as the ratio of normalized counts in a pulldown condition vs the apo-Cas13a condition. Variation between samples was regularized by subtracting the average fold-change for an off-target sequence and then dividing by the average fold-change for an on-target sequence [FC_regularized = (FC – A) / B, where A =averaged FC for off-target, and B = averaged FC for on-target). The effect of this regularization is that sequences that bind extremely poorly have a score near 0, while sequences that bind on average as well as an on-target sequence have a score near 1. This correction results in good agreement between replicates [for example, see Supp. Fig 5]. These fold-change values were used to compute Fig.s 4a. To calculate the statistical significance of these observations, we performed permutation tests. To do this, we drew a random sample of 9 points each from the two targets, and record the difference in the means between these two sub-samplings. We repeated this sampling process 100,000 times to produce an empirical distribution of the difference in means for the guide-complementary and guide-non-complementary sequences. This empirical distribution of subsampled fold changes was used to compute a p-value for observing a given difference in means (p < 0.05).

To identify highly enriched or depleted nucleotide / position combinations (Fig. 6b), we ranked sequences by fold change (relative to the apo-Cas13a control) and computed the frequencies of each nucleotide at each position for the top 500 or bottom 500 sequence entries in this ranked list. To verify that the observed frequencies are significant, they were compared against a null hypothesis generated by permuting the order of this ranked list (i.e. shuffling the fold-changes), using 10,000 permutations. We then used a two-tailed t-test and a p-value cut-off of p <= 0.001 to identify frequency / position combinations that were significantly enriched / depleted in the fold change ranking.

## ONLINE DATA

All sequencing reads are available through the NCBI SRA BioProject ID: PRJNA399726, and all scripts and analysis presented here is publically available at github.com/OConnell-lab-URMC/High-throughput-RNA-mismatch-profiling.

## ACKNOWLEDGEMENTS

We thank Nan Ma and Kaihong Zhou for technical assistance. We thank Dmitri Ermolenko, and members of the O’Connell and Doudna laboratories for helpful discussions. We thank Shana McDevitt and Dylan Smith for high throughput sequencing assistance and helpful advice. We thank Laura Lavery and Callison Alcott for the pDuBir plasmid and tips on in vivo biotinylation. This work used the Vincent J. Coates Genomics Sequencing Laboratory at UC Berkeley, supported by the NIH S10 OD018174 Instrumentation Grant, and the Typhoon FLA9500 scanner at the Center for RNA Biology, University of Rochester, supported by the NIH S10 OD021489-01A1 Instrumentation Grant. A.T. was supported by the HARC Center National Institutes of Health Grant P50GM82250 (N. Krogan, PI; J.A.D., co-PI). G.J.K. acknowledges support from the Howard Hughes Medical Institute. M.R.O. was supported by a C.J Martin Overseas Early Career Fellowship from the National Health and Medical Research Council of Australia. This work was supported in part by a Frontiers Science award from the Paul Allen Institute to J.A.D, the National Science Foundation (MCB-1244557 to J.A.D.) and the National Institutes of Health Center for RNA Systems Biology Grant P50-GM102706 (J. Cate, P.I.; J.A.D., co-PI). J.A.D is an Investigator of the Howard Hughes Medical Institute. J.A.D. is a co-founder of Caribou Biosciences, Inc., Editas Medicine, and Intellia Therapeutics.

## AUTHOR CONTRIBUTIONS

M.R.O, A.T, A.E-S designed the project with input from G.J.K and J.A.D.

M.R.O, A.T, A-E.S carried out experiments. A.T. developed the bioinformatics pipeline, and analyzed all the sequencing data with assistance from M.R.O. M.R.O drafted the manuscript with assistance from A.T., and all authors reviewed and edited the manuscript.

---

**Figure S1:**
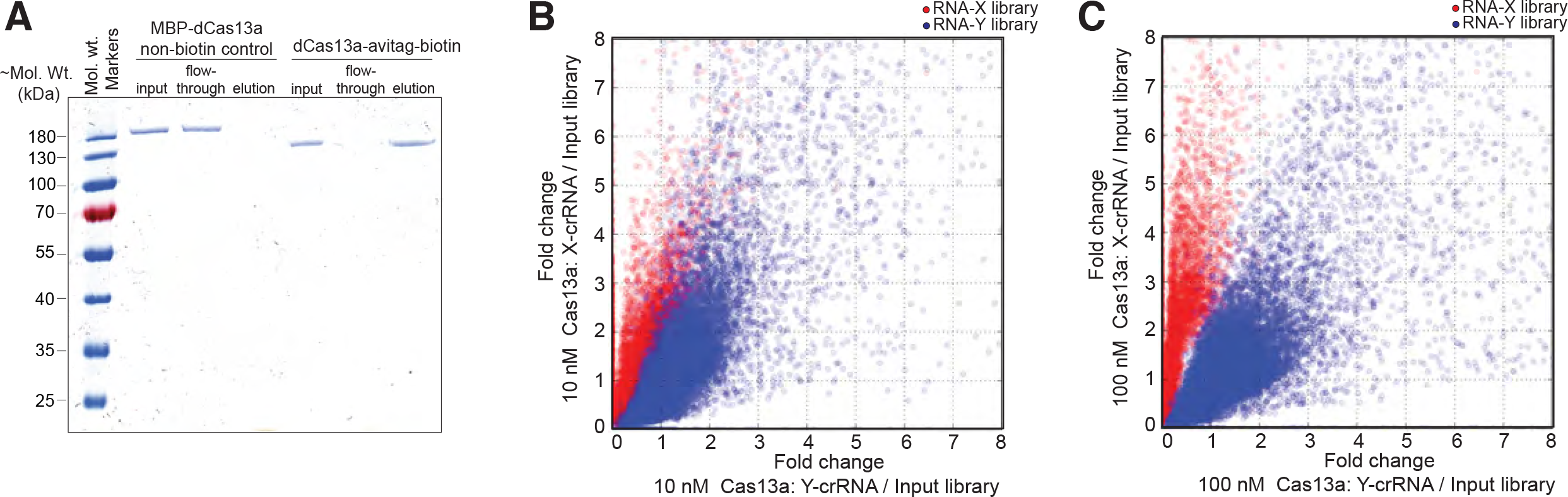
Related to Figure 1. Controls for the development of high-throughput RNA mismatch profiling to determine activator-RNA binding preferences for Lbu-Cas13a. (A) Streptavidin pulldown experiment to confirm Lbu-dCas13a-avitag biotinylation. Streptavidin bead binding assay with biotinylated Lbu-dCas13a-avitag or non-biotinylated MBP-Lbu-Cas13a and streptavidin magnetic beads. 8.5 pmol of biotinylated Lbu-dCas13a-avitag or non-biotinylated MBP-Lbu-Cas13a were separately mixed with 12 μL streptavidin magnetic beads and incubated in modified processing buffer at 37C for 1 h, followed by three washes with modified processing buffer and elution in boiling SDS-PAGE loading buffer. Input, incubation flow-through & elution for each sample were analyzed using SDS-PAGE. Biotinylated Lbu-dCas13a-avitag only remains specifically bound to the beads. A scatter plot displaying fold-change in abundance of each RNA-X and –Y library members between samples with(B) 10 nM crRNA-bound Cas13a (vs. input library) or (C) 10 nM crRNA-bound Cas13a (vs. input library) programmed to target RNA-X or -Y ssRNAs.

**Figure S2:**
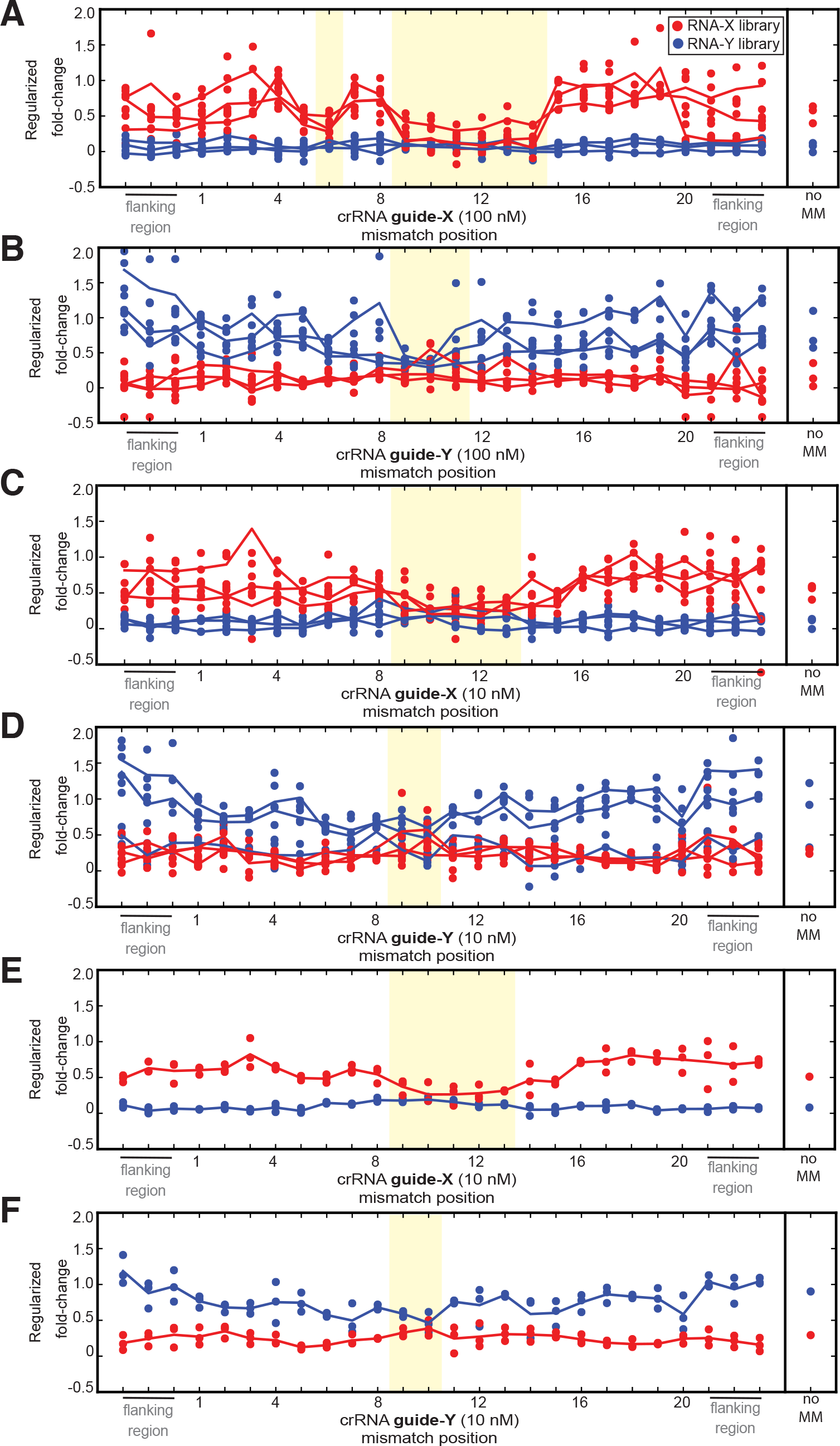
Related to Figure 2. Additional dCas13a high-throughput RNA mismatch profiling data. Regularized fold-change enrichment analysis (relative to Apo-dCas13a) for activator-RNA members of both libraries that contain only a single mismatch across the shown spacer-targeting and flanking regions. Plotting all individual points for (A) targeting X activator-RNA library with 100 nM dCas13a: crRNA-X, (B) targeting Y activator-RNA library with 100 nM dCas13a: crRNA-Y,(C) targeting X activator-RNA library with 10 nM dCas13a: crRNA-X, and (D) targeting Y activator-RNA library with 10 nM dCas13a: crRNA-Y. In each plot, each individual point in a position represents the fold-change value for each unique mismatch combination possible in that position (from n=3 experiments), The solid line represents as an average fold-enrichment (of all mismatches combinations) in that position for each experiment. Plotting average points for (E) targeting X activator-RNA library with 10 nM dCas13a: crRNA-X, and (F) targeting Y activator-RNA library with 10 nM dCas13a: crRNA-Y. In each plot, each individual point in a position represents the average regularized fold-change value for each unique mismatch combination possible in that position (from n=3). The solid line represents as an average fold-enrichment (of all mismatches combinations) in that position. The fold-change enrichment of the perfectly complementary (noMM) targets in each experiment are show on the right of each plot. The fold-change enrichment of the perfectly complementary (noMM) targets in each experiment are show on the right of each plot. Positions shaded in yellow indicate where statistically significant (p < 0.05) lack of enrichment between guide-complementary and guide-non-complementary sequences was observed. See Methods for regularization and permutation test procedure.

**Figure S3:**
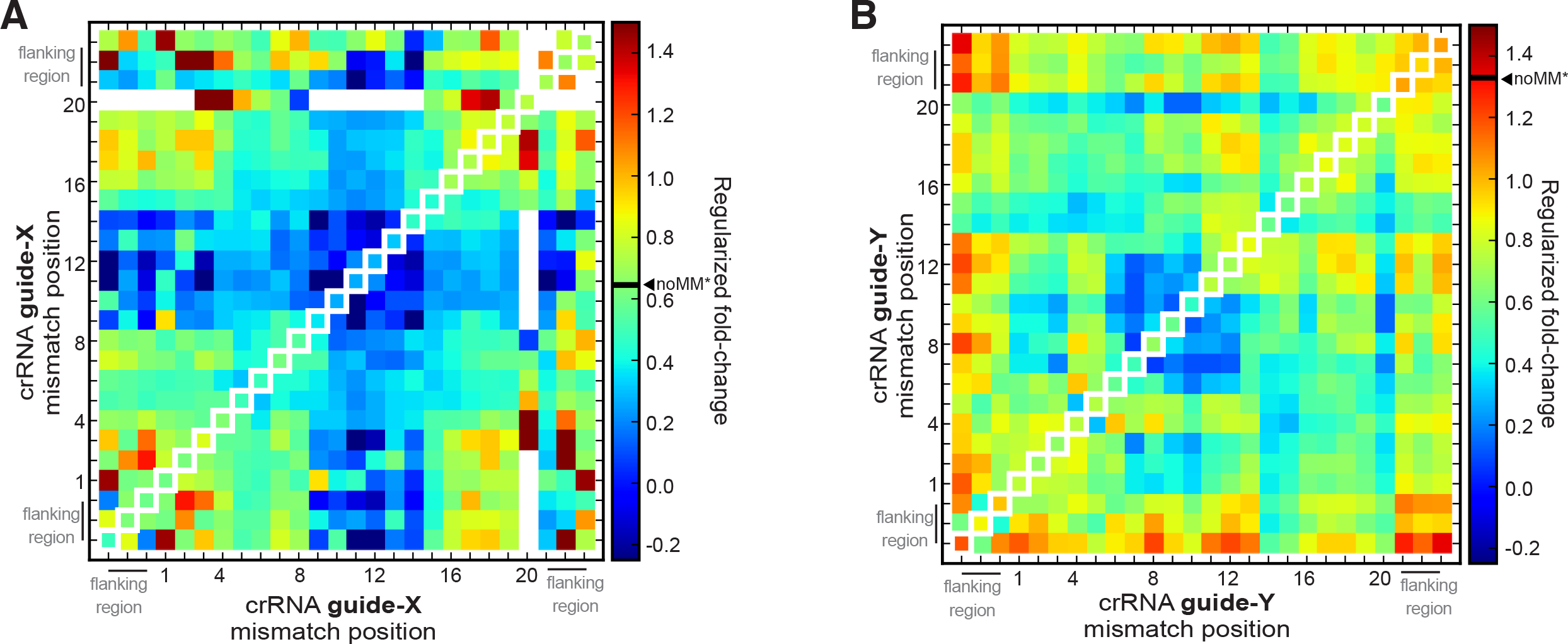
Related to Figure 2. Additional data demonstrating that dCas13a high-throughput RNA mismatch profiling confirms binding mismatch sensitivity hotspots for double-mismatched activator-RNAs. Regularized fold-change enrichment analysis (relative to Apo-dCas13a) for activator-RNA members of both libraries for all pairs of mismatches across the shown spacer-targeting and flanking regions when (A) targeting X activator-RNA library with 10 nM dCas13a: crRNA-X, and (B) targeting Y activator-RNA library with 10 nM dCas13a: crRNA-Y. For comparison, single mismatch data is plotted along the diagonal axis and the fold-change value for a perfectly complementary target is indicated on the heat-map scale bar (noMM*).

**Figure S4:**
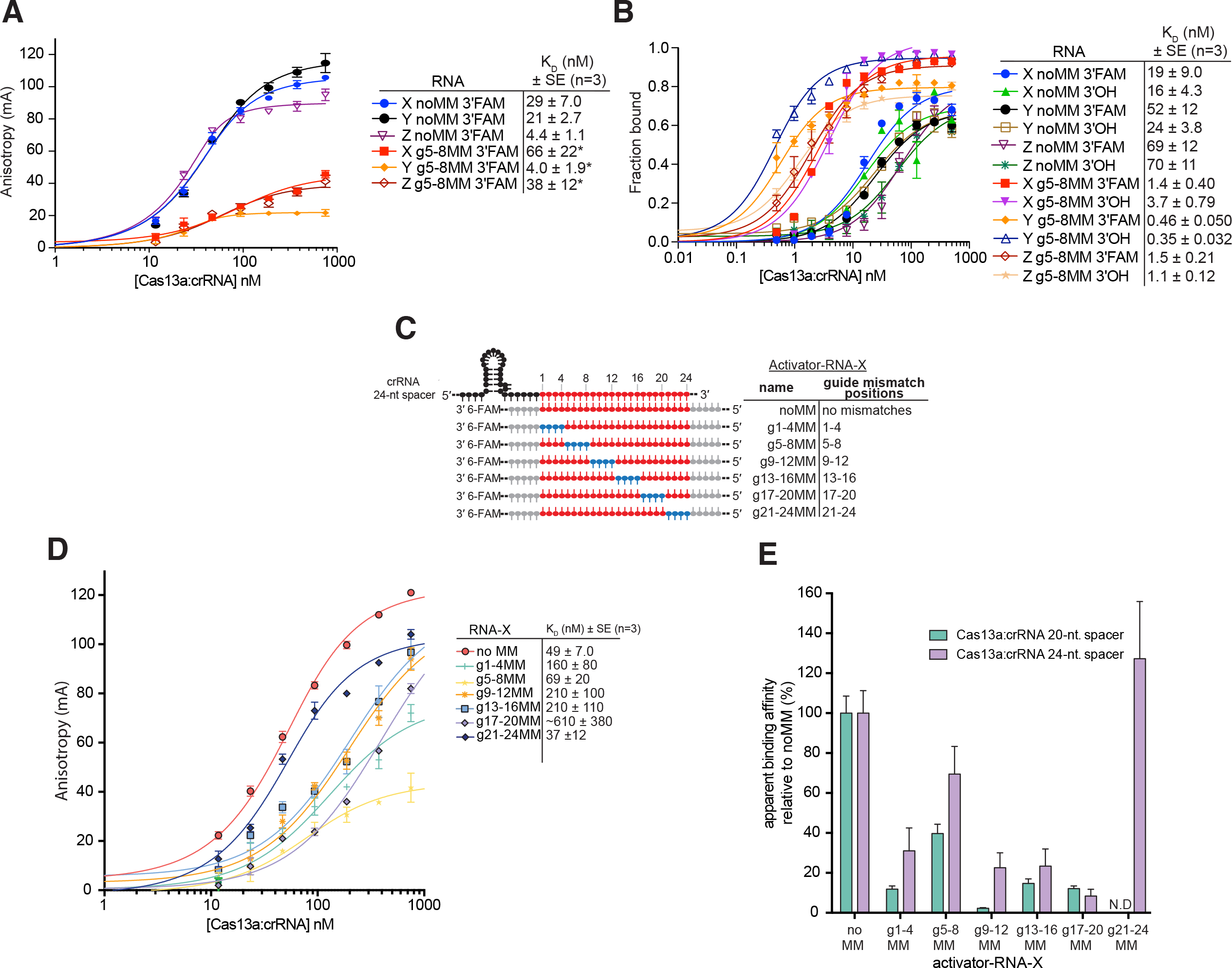
Related to Figure 3 & 4. Additional activator-RNA binding affinity experiments to validate g5-8MM behavior and additional binding affinity experiments using a 24-nt. crRNA-spacer. (A) Fluorescence anisotropy binding curves for the interaction between Lbu-dCas13a: crRNA (X, Y or Z) and activator-RNAs (X, Y or Z) with noMM or g5-8MM mismatches. Binding data fits (see Methods) shown as solid lines. Error-bars represent the s.d of the fraction bound from three independent experiments. Dissociation constants (KD) and their associated standard errors (SE) from three independent experiments are shown. An asterisk (*) indicates KD measurements for g5-8MM targets that are very likely inaccurate due to very low change in fluorescent anisotropy between unbound and bound activator RNA species. (B) Filter binding curves for the interaction between Lbu-dCas13a: crRNA (X, Y or Z) and activator-RNAs (X, Y or Z) with noMM or g5-8MM mismatches. Activator-RNAs are all labeled with a 5’-32P, and either a free 3’-OH or a 3’-6FAM fluorophore, and were detected using phosphor-imaging. Binding data fits (see Methods) shown as solid lines. Error-bars represent the s.d of the fraction bound from three independent experiments. Dissociation constants (KD) and their associated standard errors (SE) from three independent experiments are shown. (C) Schematic of the Lbu-Cas13a crRNA-24-nt.-spacer: activator-RNA interaction highlighting the different activator-RNAs tested in this study. (D) Fluorescence anisotropy binding curves for the interaction between Lbu-dCas13a: crRNA-X (with a 24-nt. spacer) and X activator-RNAs depicted in (C). Binding data fits (see Methods) shown as solid lines. Error-bars represent the s.d of the anisotropy from three independent experiments. Dissociation constants (KD) and their associated standard errors (SE) from three independent experiments are shown. (E) Data from (D) normalized as a percentage binding affinity relative to the affinity to a perfectly complementary activator-RNA X (no MM).

**Figure S5:**
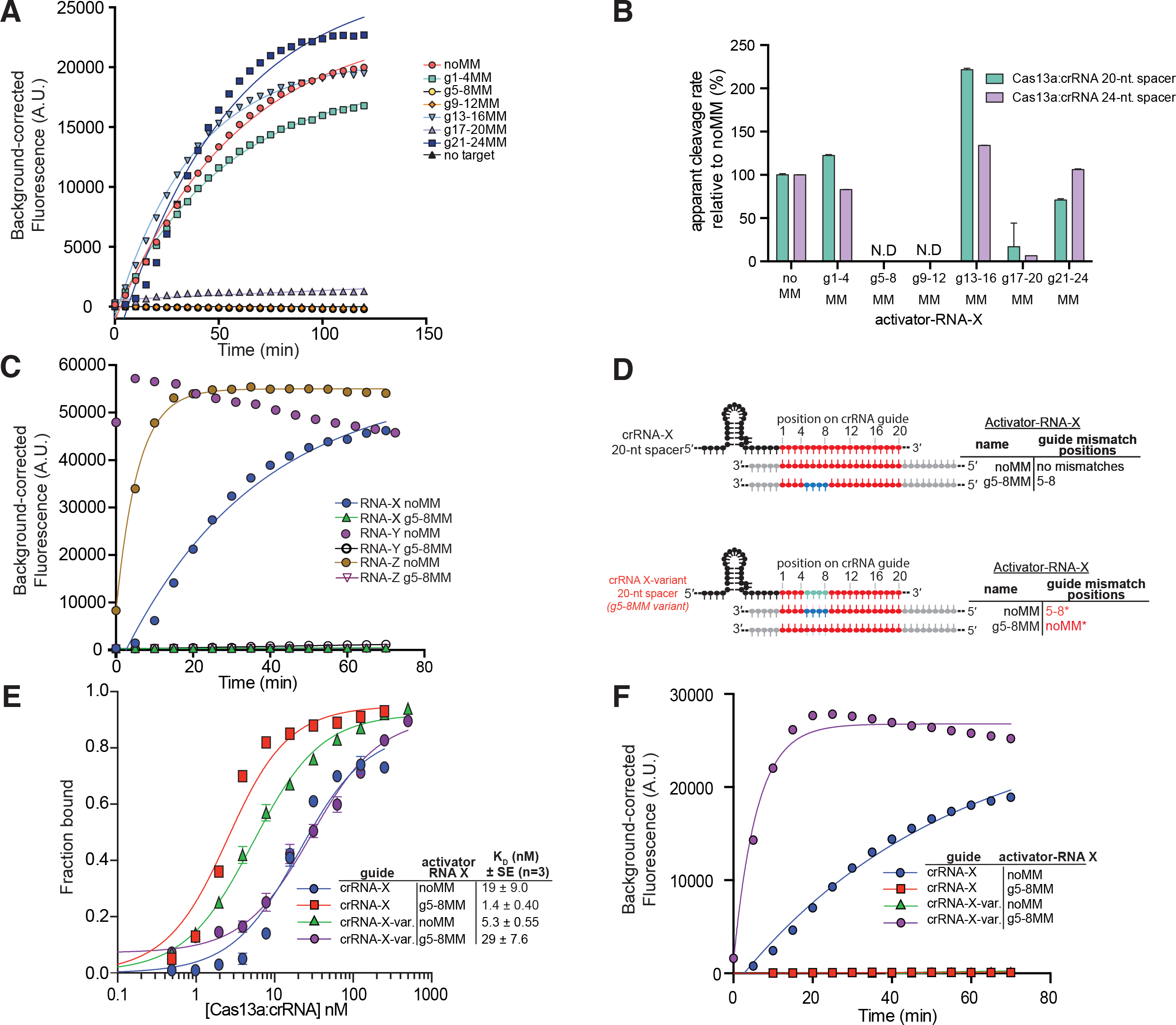
Related to Figure 4. Additional HEPN-nuclease activation experiments with a 24-nt. crRNA-spacer, and additional data showing g5-8MM abolish Lbu-dCas13a HEPN-nuclease activation but not binding. (A) representative time course of background corrected fluorescence measurements generated by Lbu-Cas13a: crRNA-X (with a 24-nt. spacer) activation by the addition of 100 pM X activator-RNAs with either zero (noMM) or four consecutive mismatches across the crRNA-spacer region. Quantified data were fitted with single-exponential decays (solid line) with calculated pseudo-first-order rates constants (k_obs_) (mean ± s.d., n =3) as follows: noMM 0.025 ± 0.001 min^-1^, g1-4MM 0.021 ± 0.001 min^-1^, g13-16MM 0.034 ± 0.001 min^-1^, g17-20MM 0.002 ± 0.010 min^-1^, g21-24MM 0.027 ± 0.003 min^-1^ while g5-8MM and g9-12 were not fit. (B) Data from (A) normalized as a percent cleavage rate relative to a perfectly complementary X activator-RNA (noMM) with error bars representing the normalized s.d from three independent experiments. (C) representative time courses of background corrected fluorescence measurements generated by Lbu-Cas13a activation by the addition of 100 pM activator-RNAs with either zero (noMM) or four consecutive mismatches in positions 5-8 of the crRNA-spacer (g5-8MM). Three different RNA sequences (X, Y & Z) and their corresponding crRNAs were tested. Exponential fits are shown as solid lines, except no fit is shown for RNA-Y noMM, as this reaction when to completion before accurate measurements could be taken. (D) Schematic of the Lbu-Cas13a crRNA: ssRNA activator interaction highlighting the position of the mismatches between noMM and g5-8MM RNAs with standard crRNA-X (std) and crRNA-X-variant (X-var.; with mismatches in positions 5-8 of the crRNA). (E) Filter binding curves for the interaction between Lbu-dCas13a: crRNA (X, or X-var.) and activator-RNA-X with noMM or g5-8MM mismatches. Activator-RNAs are all labeled with a 5’-32P, contain a 3’-6FAM fluorophore, and were detected using phosphor-imaging. Binding data fits (see Methods) shown as solid lines. Error-bars represent the s.d of the fraction bound from three independent experiments. Dissociation constants (KD) and their associated standard errors (SE) from three independent experiments are shown. (F) representative time courses of background corrected fluorescence measurements generated by crRNA-X std- or crRNA-X-variant-loaded Lbu-Cas13a activated by the addition of 100 pM activator-RNA-X noMM or RNA-X g5-8MM. Exponential fits are shown as solid lines.

**Figure S6:**
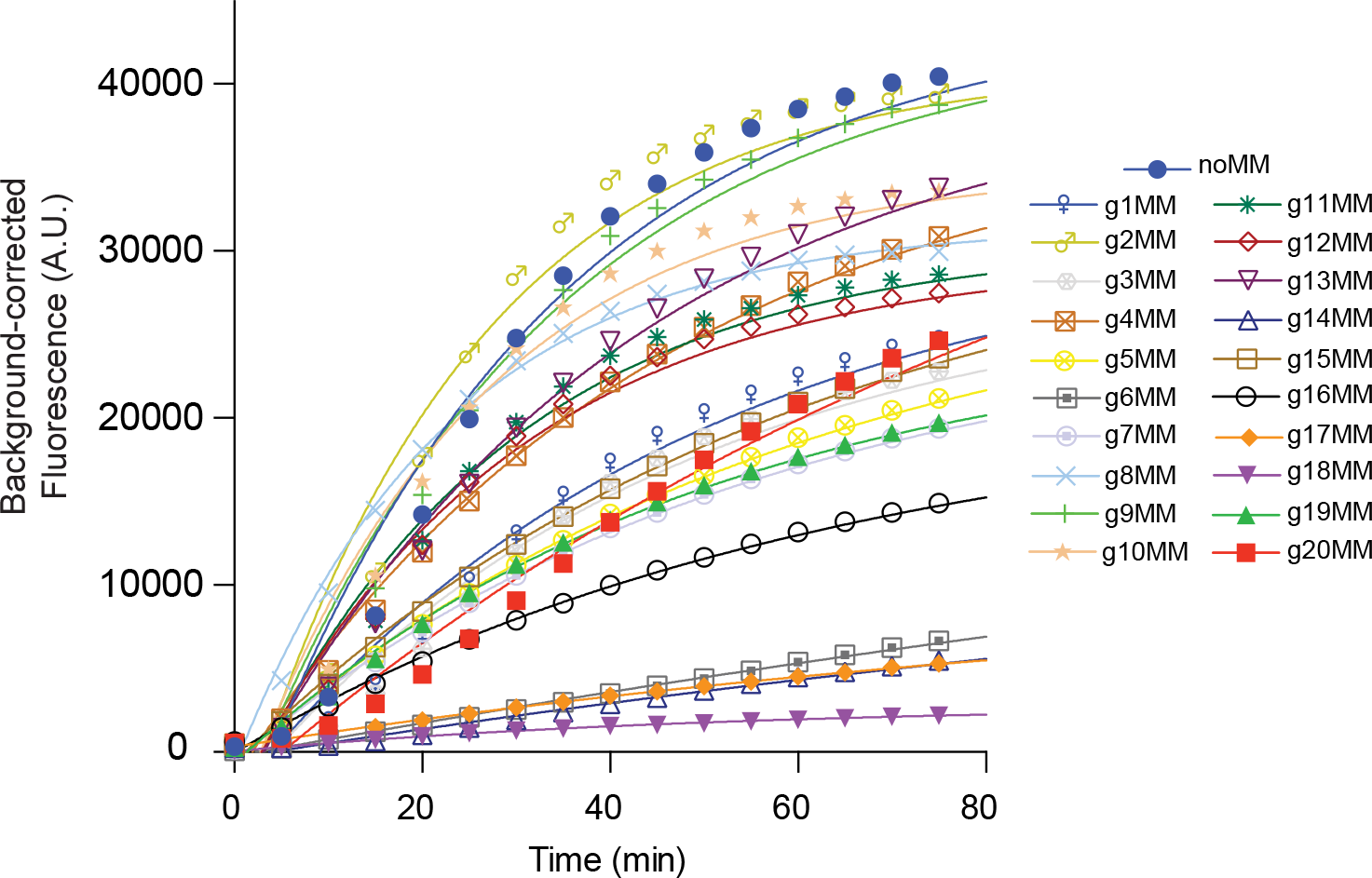
Related to Figure 5. Single mismatch HEPN-nuclease activation time course data. (A) representative time courses of background corrected fluorescence measurements generated by Lbu-Cas13a: crRNA-X activation by the addition of 100 pM λ4 activator-RNAs with either zero or single mismatches within positions 1-20 of crRNA-spacer targeting region. Exponential fits are shown as solid lines. Quantified data were fitted with single-exponential decays (solid line) with calculated pseudo-first-order rates constants (k_obs_) (mean ± s.d., n =3) as follows: noMM 0.031 ± 0.003 min^-1^, g1MM 0.022 ± 0.002 min^-1^, g2MM 0.044 ± 0.008 min^-1^, g3MM 0.018 ± 0.001 min^-1^, g4MM 0.027 ± 0.002 min^-1^, g5MM 0.023 ± 0.001 min^-1^, g6MM 0.0058 ± 0.0002 min^-1^, g7MM 0.013 ± 0.0005 min^-1^, g8MM 0.0124 ± 0.0005 min^-1^, g9MM 0.020 ± 0.001 min^-1^, g10MM 0.06 ± 0.02 min^-1^, g11MM 0.043 ± 0.006 min^-1^, g12MM 0.048 ± 0.008 min^-1^, g13MM 0.038 ± 0.005 min^-1^, g14MM 0.017 ± 0.001 min^-1^, g15MM 0.047 ± 0.007 min^-1^, g16MM 0.017 ± 0.001 min^-1^, g17MM 0.009 ± 0.0002 min^-1^, g18MM 0.018 ± 0.001 min^-1^, g19MM 0.021 ± 0.001 min^-1^, g20MM 0.010 ± 0.0003 min^-1^.

**Table S1:**
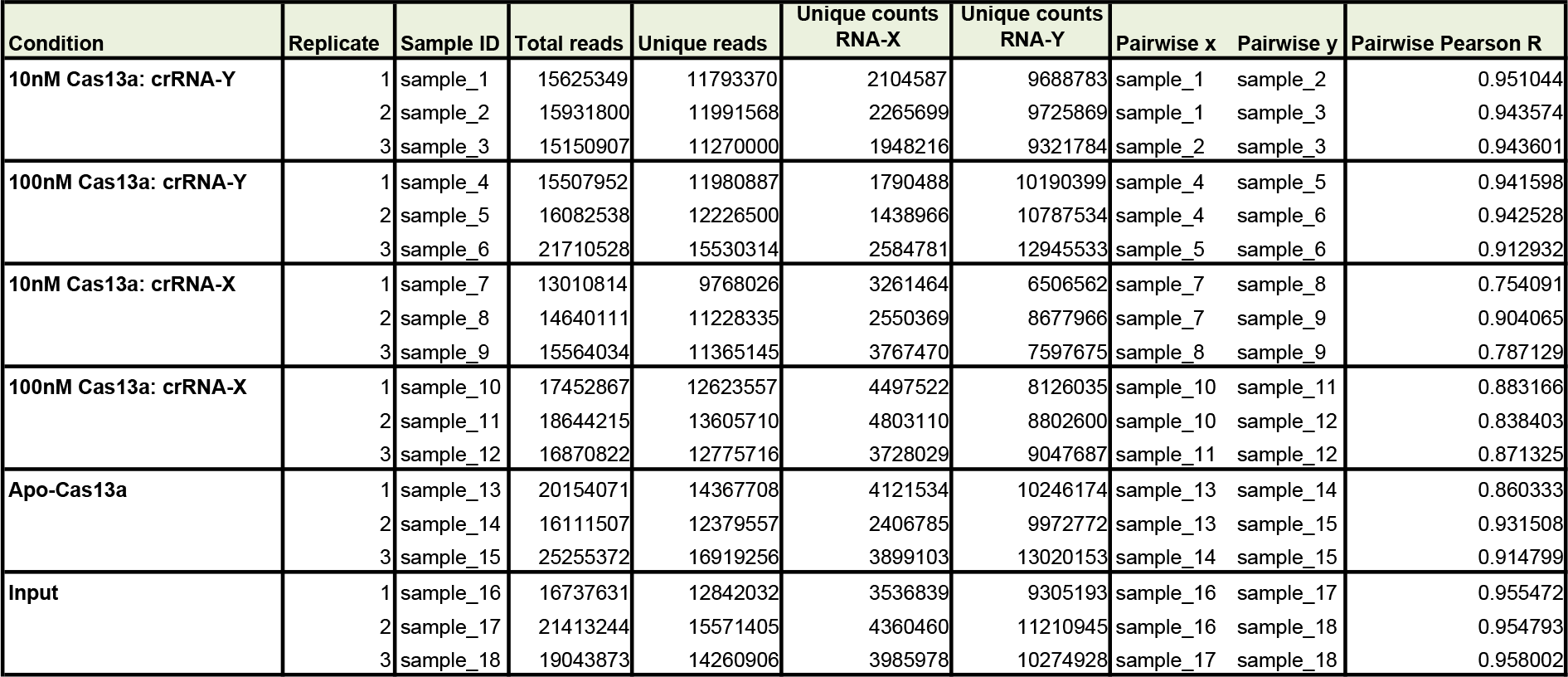
Illumina sequencing statistics for each HT-RBP sample

**Table S2:**
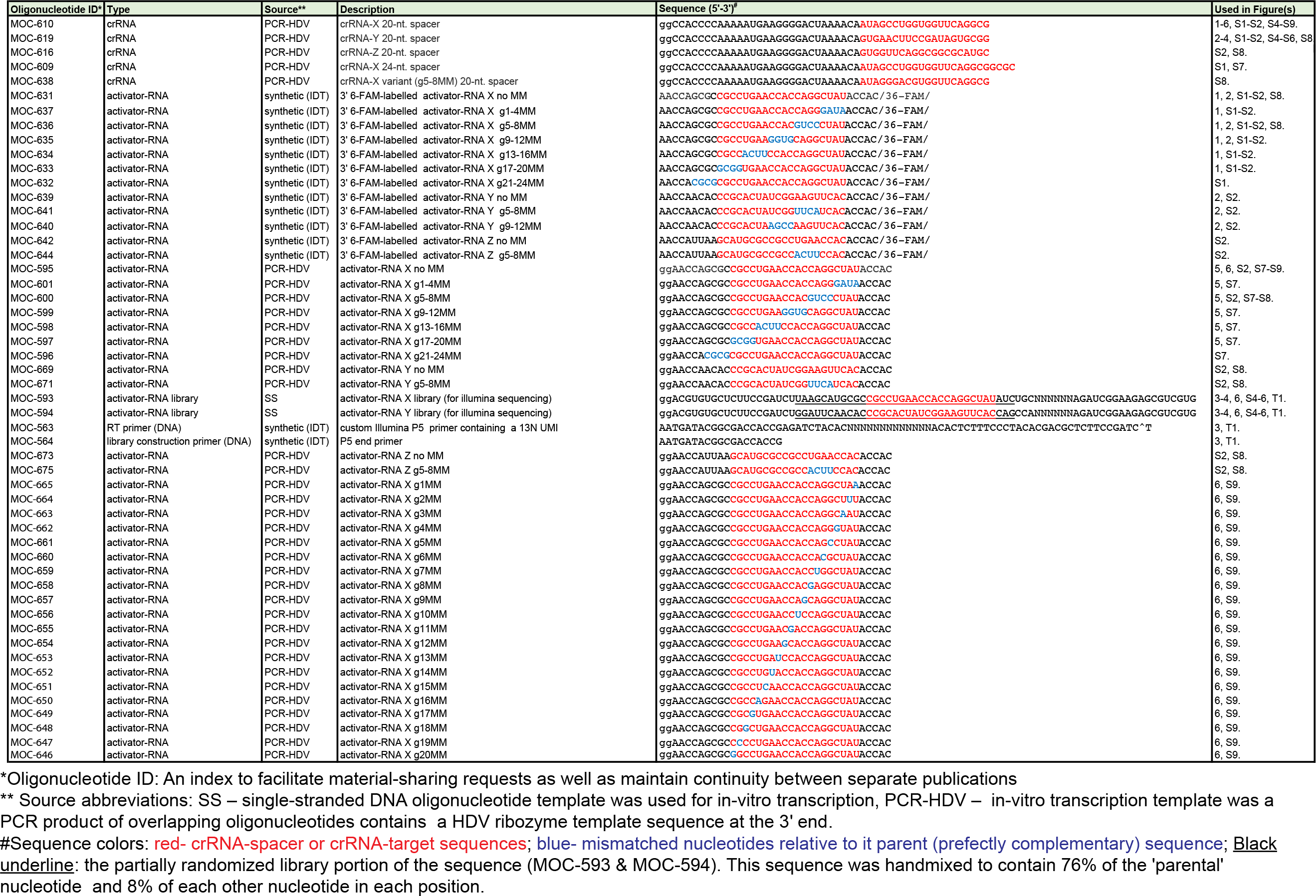
Oligonucleotide sequences used in this study

